# Female germline expression of OVO transcription factor bridges *Drosophila* generations

**DOI:** 10.1101/2023.08.25.554887

**Authors:** Leif Benner, Savannah Muron, Brian Oliver

**Affiliations:** Section of Developmental Genomics, Laboratory of Biochemistry and Genetics, National Institute of Diabetes and Digestive and Kidney Diseases, National Institutes of Health, Bethesda, MD, USA; Department of Biology, Johns Hopkins University, Baltimore, MD, USA

**Keywords:** *ovo*, zinc-finger, germ cell, oogenesis, *Drosophila melanogaster*, pole cell, embryogenesis

## Abstract

OVO is required for karyotypically female germ cell viability but has no known function in the male germline in Drosophila. *ovo* is autoregulated by two antagonistic isoforms, OVO-A and OVO-B. All *ovo^-^* alleles were created as partial revertants of the antimorphic *ovo^D1^*allele. Creation of new targeted alleles in an *ovo^+^* background indicated that disrupting the germline-specific exon extension of *ovo-B* leads to an arrested egg chamber phenotype, rather than germ cell death. RNA-seq analysis, including >1K full length cDNAs, indicates that *ovo* utilizes a number of unannotated splice variations in the extended exon and a minor population of *ovo-B* transcripts utilizes an alternative splice. This indicates that classical *ovo* alleles such as *ovo^D1rv23^*, are not truly null for *ovo*, and are likely to be weak antimorphs. To generate bonafide nulls, we deleted the *ovo-A* and *ovo-B* promoters showing that only *ovo-B* is required for female germ cell viability and there is an early and polyphasic developmental requirement for *ovo-B* in the female germline. To visualize OVO expression and localization, we endogenously tagged *ovo* and found nuclear OVO in all differentiating female germ cells throughout oogenesis in adults. We also found that OVO is maternally deposited into the embryo, where it showed nuclear localization in newly formed pole cells. Maternal OVO persisted in embryonic germ cells until zygotic OVO expression was detectable, suggesting that there is continuous nuclear OVO expression in the female germline in the transition from one generation to the next.

**Article Summary:** *ovo* has long been considered to be at the top of the female germline sex determination pathway. We utilized updated genetic methods to determine OVO expression, localization, and requirement in the embryonic and adult germline. Our results indicate that OVO is always present, and likely required, in the Drosophila female germline.

## Introduction

Sexual reproduction is a fundamental aspect to the continuity of life. Sexually reproducing organisms contain germ cells, which eventually differentiate into sex-specific gametes. Germ cells are generally specified early on in development, and organisms have different mechanisms for inducing germ cell formation. Once the germline is established and development ensues, a genetic program is established instructing the germline to differentiate into the appropriate sex-specific gamete, an egg or sperm. Sperm are essentially motile packets of DNA, while eggs in contrast, contain the RNAs and proteins needed for early development after fertilization. These two gametes are incredibly diverse, and it is still puzzling as to how germ cells, that initially appear to be the same in the sexes, can differentiate into drastically different cell types. We currently have little insight into how these sex-specific genetic programs are established to give rise to an egg or sperm, which is one of the most conserved and fundamentally important processes in biology.

The adult *Drosophila* reproductive structures house germline stem cells that divide into daughter cells that eventually differentiate into either an egg or sperm, a process called oogenesis and spermatogenesis, respectively. Both male and female germline stem cells reside in special microenvironments, where they receive signals from somatic support cells (Fuller and Spradling 2007). The sexual identity of germ cells are influenced by both the karyotype of the germ cell itself, as well as the karyotype of the surrounding somatic support cells. Incompatibility of these signals results in either germ cell death or a tumor phenotype, and thus leads to an inability to properly differentiate into the correct gamete and sterility (Steinmann-Zwicky, Schmid, and Nöthiger 1989; Schüpbach 1985, 1982; Nöthiger et al. 1989). The autonomous and non-autonomous signals influencing germline sexual identity has been a major research focus in the field. A number of important genes and pathways have been established, yet we currently have a poor understanding of these genetic programs and how they contribute to the proper differentiation of gametes. This is an important question to elucidate if we are to understand sexual reproduction and early embryonic development across species.

One candidate for the establishment of the female specific germline genetic network is the transcription factor OVO. The expression of *ovo* is autonomously influenced by the X karyotype of the germline (XX = female, X = male). XX germ cells strongly express *ovo* while single X germ cells express *ovo* at low levels (B. Oliver et al. 1994; Hinson and Nagoshi 1999; Andrews and Oliver 2002). *Drosophila* female germ cells lacking OVO^+^ activity are not viable, while male germ cells are unaffected, and it is therefore an essential protein for the proper viability and differentiation of the female germline (B. Oliver, Perrimon, and Mahowald 1987). OVO has been shown to positively regulate the transcription of itself and a downstream target gene *ovarian tumor* (*otu*) (Lü et al. 1998; Lü and Oliver 2001; Andrews et al. 2000; Bielinska et al. 2005). *otu* is also required for proper female germ cell differentiation, with *otu^-^*germ cells having phenotypes ranging from germ cell death to tumorous ovaries (Pauli, Oliver, and Mahowald 1993; Bishop and King 1984). The presence of OTU is needed for the proper splicing and expression of *Sex lethal* (*Sxl*) (Bopp et al. 1993; B. Oliver, Kim, and Baker 1993; Pauli, Oliver, and Mahowald 1993). *Sxl* has been shown to be both necessary and sufficient for germline feminization, where loss of function leads to a similar ovarian tumor phenotype seen with hypomorphic *otu* alleles (Hashiyama, Hayashi, and Kobayashi 2011; Schüpbach 1985). Loss of function of *ovo*, *otu*, or *Sxl* has no effect on male germline differentiation, so therefore the *ovo*→*otu*→*Sxl* pathway is considered to be the core female germline sex-determination pathway (Brian Oliver 2002).

The *ovo* locus, which shares the same reading frame with *shavenbaby* (*svb*), expresses multiple transcripts, all of which encode four C_2_H_2_ zinc fingers at the C-terminal domain with variable N-termini. SVB, the somatic version of OVO, is an essential protein for embryonic and first instar development (B. Oliver, Perrimon, and Mahowald 1987). *svb* transcripts arise from an upstream promoter expressed in various somatic cells and encodes a protein that switches from a transcriptionally repressive larger isoform to a transcriptionally activating smaller isoform via *pri* dependent proteolytic cleavage (T. Kondo et al. 2010). In contrast, *ovo* utilizes two alternative promoters that are expressed exclusively in the germline (Andrews and Oliver 2002). An upstream promoter encodes the transcriptionally repressive OVO-A isoform that contains the same repressive protein domains as the repressive SVB isoform. While a downstream promoter encodes the transcriptional activator OVO-B, which lacks the repressive protein domain of OVO-A, and at the protein level is similar to the transcriptional activating SVB isoform (Andrews et al. 2000). The five antimorphic *ovo^D^* alleles (three from a screen and two engineered) illustrate this dual activity between OVO-A and OVO-B (Busson et al. 1983; Andrews, Levenson, and Oliver 1998). The three alleles from a dominant female sterility screen all create in-frame AUGs that add the repressive protein domains of OVO-A to the N-terminus of the OVO-B open reading frame (Busson et al. 1983; Mével-Ninio et al. 1996). Design of new in-frame AUGs in the *ovo-B* 5’-UTR or over-expression of OVO-A encoding transgenes are also dominant antimorphs, clearly showing the deleterious effect of these polypeptide sequences on the female germline (Andrews, Levenson, and Oliver 1998). This strong *ovo^D1^* phenotype has been widely exploited in mitotic recombination experiments to assay for the maternal effects of lethal alleles in germline versus somatic clones (Perrimon 1984). The two most widely used amorphs of *ovo* (*ovo^D1rv22^* and *ovo^D1rv23^*) are both revertants of *ovo^D1^* and thus double mutants (B. Oliver, Perrimon, and Mahowald 1987). In one case, by a *Gypsy* insertion near the *ovo-B* promoter (Mével-Ninio, Mariol, and Gans 1989; Garfinkel, Lohe, and Mahowald 1992) and in the other, a *Stalker2* insertion inside an alternatively spliced extended exon 2 (Salles et al. 2002). This alternative exon 2 extension is the only other annotated difference between the OVO-A and OVO-B isoforms, where *ovo-A* transcripts have been annotated to use an upstream alternative splice in exon 2 to exon 3 (like *svb*). *ovo-B* is annotated to use a downstream alternative splice site from exon 2 into exon 3 and therefore contains an extended exon 2 (G. dos Santos et al. 2015; Gramates et al. 2022).

Herein, we utilize updated genetic methods to investigate OVO localization, expression, and requirements in the adult and embryonic *Drosophila* germline. Newly generated *ovo* alleles and small and long read RNA-seq analysis at the *ovo* locus suggest that our current understanding of the complex genetics of the *ovo* locus are likely incorrect and we suggest a more representative *ovo* genetic model. We show that OVO is required in the female germline and is highly expressed, localizing to differentiating germ cell nuclei throughout oogenesis. OVO is maternally deposited, where maternal OVO persists in male and female germ cells until late into embryogenesis where it overlaps with the beginning of zygotic OVO expression, independent of sex. We show that at the end of embryogenesis, there seems to be an early requirement for OVO in the female germline, whereas if this requirement for OVO is not met, adult females lack germ cells and are sterile. There appears to be a second function of *ovo* as oogenesis begins in the adult ovary. Altogether, our study suggests that OVO is an important female-specific germline transcription factor that is eternally present and likely to be continuously required for female germ cell viability and differentiation.

## Methods

All reagents used in this study can be found in the FlyBase recommended supplementary ART Table (Table S1).

### Fly Husbandry, Transgenesis, and CRISPR/Cas9

Flies were kept under constant light and all crosses were conducted at 25°C with 65% relative humidity. Flies were fed on premade flyfood (D20302Y) from Archon Scientific (Durham, NC). All *attB* plasmids were integrated into *P{CaryP}Msp300^attP40^* with *P{nanos-phiC31\int.NLS}X* as the integration source (Bischof et al. 2007; Markstein et al. 2008). *y^1^ sc* v^1^ sev^21^*; *P{nanos-Cas9.R}attP40* flies (Ren et al. 2013) were injected with 500 ng/μL donor plasmid and 150 ng/μL gRNA plasmid to generate the *ovo^ovo-GAL4^* allele. *y^1^ w^1118^*; *P{nanos-Cas9.K}attP2* flies (S. Kondo and Ueda 2013) were injected with 500 ng/μL donor plasmid and 150 ng/μL gRNA plasmid to generate all other *ovo* alleles used in this study. To generate *ovo^ΔAP-Nterm-3xFHA^*, *ovo^Nterm-3xFHA^*/*Y*; *P{nanos-Cas9.R}attP40*/*P{U6:3-gRNA^ovo-ΔAP^}attP40* males were crossed to *C(1)DX*/*Y* females and the resulting F1 male progeny were individually backcrossed to *C(1)DX*/*Y* females. Subsequent F2 males were PCR screened with ovo-deltaAP-F/ovo-deltaAP-R primers and Sanger sequenced with ovo-deltaAP-F primer in order to recover *ovo^ΔAP-Nterm-3xFHA^*. *ovo^ΔAP-Cterm-3xFHA^*was generated with the same method, but with *ovo^Cterm-3xFHA^*/*Y* males instead. To generate *ovo^ext–^* flies, males with the genotype *y^1^ cho^2^ v^24^ f^1^ P{ry+t7.2=neoFRT}19A*; *P{nanos-Cas9.R}attP40*/*P{U6:3-gRNA^ovo-ext–^}attP40*, were crossed to *C(1)DX*/*Y* females and the resulting F1 male progeny were individually crossed to *ovo^ΔBP^*/*FM7c* females. Resulting F2 female progeny with the genotype *ovo^ΔBP^*/*y^1^ cho^2^ v^24^ f^1^ P{ry+t7.2=neoFRT}19A* were scored for fertility, and if sterile, indicating a mutation at the *ovo* locus, sister *y^1^ cho^2^ ovo* v^24^ f^1^ P{ry+t7.2=neoFRT}19A*/*FM7c* females were crossed to *FM7c*/*Y* males. Subsequent *y^1^ cho^2^ ovo* v^24^ f^1^ P{ry+t7.2=neoFRT}19A*/Y males were Sanger sequenced with ovo-ext-F/ovo-ext-R primers and one line, with a single nucleotide deletion (G) at position 2,070 in the *ovo-RA* reading frame was kept and named *ovo^ext–^*. This frameshift mutation results in an early stop codon 15 codons away within the *ovo-RA* reading frame before the shared C_2_H_2_ zinc-fingers of *ovo*.

Screening for CRISPR/Cas9 generated alleles were as follows. N-terminally tagged alleles of OVO-A were PCR screened and Sanger sequenced with ovo-A-F/ovo-A-R. N-terminally tagged alleles of OVO-B/OVO-A were PCR screened and Sanger sequenced with ovo-B-F/ovo-B-R. C-terminally tagged alleles of OVO were PCR screened and Sanger sequenced with ovo-C-F/ovo-C-R. GAL4 insertion in *ovo^ovo-GAL4^* were PCR screened and Sanger sequenced with ovo-GAL4-F/ovo-GAL4-R. All injections were completed by BestGene (Chino Hills, CA). Sanger sequencing was completed by GENEWIZ (South Plainfield, NJ).

To generate *ovo^ΔAP^* flies, females with the genotype *ovo^ΔAP-FRT-DsRed-FRT^*/*FM7c*; *P{hs-FLPG5.PEST.Opt}attP40*/+ (Nern, Pfeiffer, and Rubin 2015) were heat shocked for 75 minutes at 37°C two days after egg laying. Females were crossed out and DsRed^-^ males were individually selected and crossed out. PCR was performed with ovo-deltaAP-F/ovo-deltaAP-R primers on these males to confirm the removal of the 3xP3-DsRed-SV40 termination sequence, leaving only a single FRT sequence in place of the *ovo-A* promoter. *ovo^ΔBP^* flies were generated in the same fashion except the *ovo^ΔBP-FRT-DsRed-FRT^*allele was used and PCR primers ovo-deltaBP-F/ovo-deltaBP-R were used for screening. For heat shock induced OVO rescue experiments. *ovo^D1rv22^*/*ovo^D1rv23^*; *ovoP-FRT-2xSV40-FRT-OVO-B*/*P{hs-FLPG5.PEST.Opt}attP40* females were heatshocked for 75 minutes at 37°C on the indicated day. Females were allowed to continue to develop until adulthood and then 5 day old females were dissected and ovaries were scored for containing germ cells or being agametic.

### Plasmid Construction

Plasmids designed for *UASp* and CRISPR constructs were all novelly synthesized and cloned. In order to generate *UASp* plasmids, a *3xP3*-*DsRed*-*SV40* termination-*attB*-*Gypsy* insulator-multiple cloning site (MCS)-*Gypsy* insulator was synthesized and cloned into *pUC19* at the SphI and EcoRI restriction enzyme sites to derive *pUC19-3xP3-DsRed-attB-Gypsy-MCS*. A 10x-*UASp*-MCS-*ftz* intron-*K10* terminator (first 5x *UAS* sequences are flanked with *loxP* sequences) was synthesized and cloned into *pUC19-3xP3-DsRed-attB-Gypsy-MCS* at the XhoI and SpeI restriction enzyme sites to derive *pUC19-3xP3-DsRed-attB-Gypsy-UASp-MCS-ftz-K10*. A *Mhc16* intron-MCS-*p10* terminator sequence was synthesized and cloned into *pUC19-3xP3-DsRed-attB-Gypsy-UASp-MCS-ftz-K10* at the KpnI and SpeI restriction enzyme sites deriving *pUC19-3xP3-DsRed-attB-Gypsy-UASp-mhc16-MCS-p10*, otherwise our final *pUASp* shuttle plasmid for ΦC31 integration. In order to generate our *UASp-GFP*, a GFP cDNA sequence is preceded by a nuclear localization sequence and cloned into the *UASp* plasmid at the XbaI and FseI restriction enzyme sites. For the *pUASp-3xFHA-OVO-B^ext+^*, *pUASp-3xFHA-OVO-A*, and *pUASp-3xFHA-OVO-B^ext–^*plasmids, a *3xFlag-linker-3xHA-linker* preceding an OVO-B with exon 3 extension cDNA (corresponds to the cDNA of *ovo-RD*), OVO-A cDNA (corresponds to the cDNA of *ovo-RC*), and OVO-B cDNA without exon 3 extension (corresponds to the cDNA of *ovo-RD* with the removal of nucleotides 1725-2238), respectively, was synthesized and cloned into the *pUASp* plasmid at the XbaI and FseI restriction enzyme sites. In order to construct *ovoP-FRT-2xSV40-FRT-OVO-B*, a DNA fragment was synthesized that contained *ovo-B* regulatory sequences (−830 to the transcriptional start site), *ovo-B* 5’ UTR (1 to 429), *FRT*-2x*SV40* termination-*FRT*-FseI restriction enzyme site-Kozak sequence (AATCAAA), *ovo-B* 5’ UTR (430 to 1,312), OVO-B cDNA (*ovo-RD*), and finally the *ovo-B* 3’ UTR. This fragment was cloned into *pUC19-3xP3-DsRed-attB-Gypsy-MCS* at the KpnI and AgeI restriction enzyme sites resulting in *pUC19-3xP3-DsRed-attB-Gypsy-ovoP-FRT-2xSV40-FRT-OVO-B* (*ovoP-FRT-2xSV40-FRT-OVO-B*).

In order to generate the gRNA expressing plasmids, the *DsRed* reading frame was amplified from *pUC19-3xP3-DsRed-attB-Gypsy-MCS* with DsRed-N-F/DsRed-N-R and DsRed-C-F/DsRed-C-R primers. *pUC19-3xP3-DsRed-attB-Gypsy-MCS* was then restriction enzyme digested with BamHI and NheI. PCR products were purified with QIAquick PCR Purification Kit (Qiagen) and restriction enzyme digested *pUC19-3xP3-Dsred-attB-Gypsy-MCS* was purified with a GeneJET Gel Extraction Kit (ThermoFisher Scientific).

Purified PCR products and enzyme digested plasmid were then ligated together in a gibson reaction using NEBuilder HiFi DNA Assembly Master Mix (New England Biolabs) according to the manufacturer’s protocol resulting in *pUC19-3xP3-DsRed-attB-Gypsy-MCS-no-BbsI*. This resulting plasmid changed the ‘GAAGAC’ within the *DsRed* reading frame to ‘AAAAAC’, thus removing the BbsI restriction site. A *U6:3* promoter-*tRNA^Gly^*-MCS-gRNA scaffold-*U6:3* terminator sequence was then synthesized and cloned into *pUC19-3xP3-DsRed-attB-Gypsy-MCS-no-BbsI* at the XhoI and SpeI restriction sites resulting in *pUC19-3xP3-DsRed-attB-Gypsy-U63-gRNA*, otherwise our final gRNA shuttle vector (*pgRNA*).

To create a gRNA plasmid targeting the extended exon of the *ovo-RA* transcript and used to generate the *ovo^ext–^* allele, a gRNA^ovo-ext-1^-gRNA scaffold-*tRNA^Rice-Gly^*-gRNA^ovo-ext-2^ (Benner and Oliver 2018; Port and Bullock 2016) was synthesized and cloned into *pgRNA* at the BbsI sites creating *pgRNA-ovo-ext–*. To create a gRNA plasmid targeting the C-terminus of OVO and used to generate the *ovo^Cterm-3xFHA^*and *ovo^Cterm-GFP^* alleles, a gRNA^ovo-Cterm-1^-gRNA scaffold-*tRNA^Rice-Gly^*-gRNA^ovo-Cterm-2^-gRNA scaffold-*tRNA^Rice-Gly^*-gRNA^ovo-Cterm-3^ was synthesized and cloned into *pgRNA* at the BbsI sites creating *pgRNA-ovo-Cterm*. To create a gRNA plasmid targeting the N-terminus of OVO and used to generate the *ovo^Nterm-3xFHA^* allele, a gRNA^ovo-Nterm-1^-gRNA scaffold-*tRNA^Rice-Gly^*-gRNA^ovo-Nterm-2^ was synthesized and cloned into *pgRNA* at the BbsI sites creating *pgRNA-ovo-Nterm*. To create a gRNA plasmid targeting the N-terminus of OVO-A and used to generate the *ovo^ovo-A-3xFHA^*allele, a gRNA^ovo-A-1^-gRNA scaffold-*tRNA^Rice-Gly^*-gRNA^ovo-A-2^ was synthesized and cloned into *pgRNA* at the BbsI sites creating *pgRNA-ovo-A*. To create a gRNA plasmid targeting the *ovo-A* promoter and used to generate the *ovo^ΔAP-FRT-DsRed-FRT^* allele, a gRNA^ovo-*ΔAP-1*^-gRNA scaffold-*tRNA^Rice-Gly^*-gRNA^ovo-*ΔAP-2*^ was synthesized and cloned into *pgRNA* at the BbsI sites creating *pgRNA-ovo-ΔAP*. To create a gRNA plasmid targeting the *ovo-B* promoter and used to generate the *ovo^ΔBP-FRT-DsRed-FRT^* allele, a gRNA^ovo-*ΔBP-1*^-gRNA scaffold-*tRNA^Rice-Gly^*-gRNA^ovo-*ΔBP-2*^ was synthesized and cloned into *pgRNA* at the BbsI sites creating *pgRNA-ovo-ΔBP*. Two gRNA plasmids targeting the extended exon of OVO were created to generate the *ovo^ovo-GAL4^* allele. Phosphorylated oligos ovo-intron-gRNA-1-F/ovo-intron-gRNA-1-R and ovo-intron-gRNA-2-F/ovo-intron-gRNA-2-R were used to seal a BbsI restriction digested *pU6-BbsI-chiRNA* plasmid, generating *pU6-ovo-intron-gRNA-1* and *pU6-ovo-intron-gRNA-2*.

To generate the *ovo^Nterm-3xFHA^* allele, a donor plasmid was synthesized that contained 303 nucleotides upstream of position chrX:5065456, a *3xFlag-linker-3xHA-linker*, and 395 nucleotides downstream of position chrX:5065456, and placed into *pUC57* creating *pUC57-ovo-Nterm-3xFHA*. A G>T synonymous mutation at position chrX:5065570 and a C>A synonymous mutation at position chrX:5065636 were made in the donor plasmid. To generate the *ovo^Cterm-3xFHA^* allele, a donor plasmid was synthesized that contained 353 nucleotides upstream of position chrX:5069077, a *linker-3xHA-linker-3xFlag*, and 303 nucleotides downstream of position chrX:5069077, and placed into *pUC57* creating *pUC57-ovo-Cterm-3xFHA*. A C>G synonymous mutation at position chrX:5,069,023 and a C>A synonymous mutation at position chrX:5,069,062 were made in the donor plasmid. To generate the *ovo^Cterm-GFP^* allele, a donor plasmid was synthesized that contained 553 nucleotides upstream of position chrX:5069077, a *linker-sfGFP*, and 503 nucleotides downstream of position chrX:5069077, and placed into *pUC57* creating *pUC57-ovo-Cterm-GFP*. A C>G synonymous mutation at position chrX:5,069,023 and a C>A synonymous mutation at position chrX:5,069,062 were made in the donor plasmid. To generate the *ovo^ovo-A-3xFHA^* allele, a donor plasmid was synthesized that contained 300 nucleotides upstream of position chrX:5063846, a *3xFlag-linker-3xHA-linker*, and 338 nucleotides downstream of position chrX:5063846, and placed into *pUC57* creating *pUC57-ovo-A-3xFHA*. A C>T mutation at position chrX:5063879 and a G>T mutation at position chrX:5063889 were made in the donor plasmid. To generate the *ovo^ovo-GAL4^*allele, a donor plasmid was synthesized that contained 335 nucleotides upstream of position chrX:5067778, a *T2A-GAL4-3xSTOP*, and 294 nucleotides downstream of position chrX:5067778, and placed into *pUC57* creating *pUC57-ovo-GAL4*. A C>A synonymous mutation at position chrX:5067739 and a G>C synonymous mutation at position chrX:5067778 were made in the donor plasmid. To generate the *ovo^ΔAP-DsRed^* allele, a donor plasmid was synthesized that contained 302 nucleotides upstream of position chrX:5063725, a *FRT*-(*SV40* termination-*DsRed*-*3xP3*)antisense-*FRT*, and 309 nucleotides downstream of position chrX:5063874, and placed into *pUC57* creating *pUC57-ovo-ΔAP-DsRed*. A G>C mutation at position chrX:5063888 were made in the donor plasmid. To generate the *ovo^ΔBP-DsRed^*allele, a donor plasmid was synthesized that contained 321 nucleotides upstream of position chrX:5064211, a *FRT*-(*SV40* termination-*DsRed*-*3xP3*)antisense-*FRT*, and 306 nucleotides downstream of position chrX:5064359, and placed into *pUC57* creating *pUC57-ovo-ΔBP-DsRed*. A G>C mutation at position chrX:5064195 and a G>C mutation at position chrX:5064370 were made in the donor plasmid. All gene synthesis was completed by Genscript (Piscataway, NJ).

### Immunostaining

Adult females were collected and fed for 3-5 days before ovaries were dissected and fixed in 5.14% formaldehyde (Pierce, ThermoFisher Scientific) in phosphate buffered saline (PBS, Gibco, ThermoFisher Scientific) containing 0.1% Triton X-100 (Millipore Sigma)(PBTx) for 15 minutes. Ovaries were then washed 3 times for 5 min with PBTx followed by a blocking step in PBTx supplemented with 2% normal goat serum (NGS, Invitrogen, ThermoFisher Scientific) and 0.5% bovine serum albumin (BSA, Millipore Sigma)(BBTx) for 30 minutes. Primary antibodies were diluted to their appropriate concentrations in BBTx and ovaries were incubated in primary overnight at 4°C (rabbit anti-Vasa, 1:10K; mouse anti-α-Spectrin, 1:200; rat anti-HA, 1:100; mouse anti-BAM, 1:25; chicken anti-GFP, 1:500; mouse anti-ARM, 1:200; rabbit anti-MSL-2, 1:5K). The next day, ovaries were washed 3 times for 5 min with PBTx and incubated with appropriate secondary antibodies diluted (1:500) in BBTx at room temperature for 2 hours (Alexa Fluor goat anti-rat 488, Alexa Fluor goat anti-chicken 488, Alexa Fluor goat anti-rabbit 488, Alexa Fluor goat anti-mouse 568, Alexa Fluor goat anti-rat 568, Alexa Fluor goat anti-rabbit 633; Invitrogen,ThermoFisher Scientific). Ovaries were washed 3 times for 5 min with PBTx before incubation at room temperature in 1 μg/mL DAPI (Invitrogen, ThermoFisher Scientific) solution in PBS for 30 minutes. DAPI was then removed and ovaries were stored in PBS at 4°C until mounting. To mount, ovaries were transferred to a microscope slide before adding Ultramount Aqueous Permanent Mounting Medium (Dako, Agilent) and then coverslipped. Male testes were treated in the same manner, the only difference being that they were fixed in 4.5% paraformaldehyde in PBTx for 25 minutes. All steps were completed on a rotating nutator at room temperature unless otherwise noted.

For embryo collection, flies were put on grape agar plates (Grape Agar Powder Premix diluted in 380 mL tap water, Flystuff, Genesee Scientific) supplemented with yeast paste (dry yeast from *Saccharomyces cerevisiae* resuspended in tap water). The cage was kept at 25°C to lay for 1 hour, then the grape agar plate was switched from the cage and the embryos were left at 25°C for a cumulative 2 hours for stage 4 collection, 3 hours and 50 minutes for stage 8 collection, 7.5 hours for stage 12 collection, and 12 hours for stage 15 collection. Embryos were then collected from the plate and washed 3 times with tap water. The embryos were dechorionated using a solution of 50% bleach (Clorox germicidal bleach, The Clorox Company) in tap water and gently agitated for 2 minutes. The embryos were then washed another 3 times in tap water and immediately fixed for 20 minutes at room temperature by transferring them to a tube containing 200 uL of 4% formaldehyde in PBS and 800 uL of heptane (Millipore Sigma). After fixing, the aqueous formaldehyde layer was removed and then 700 uL of methanol (ThermoFisher Scientific) was added and vortexed for one minute. All solution was removed from the embryos and they were then washed 3 times with methanol. Embryos were then stored in methanol at −20°C until staining.

Before primary staining, embryos were rehydrated using different concentrations of methanol and 0.1% Tween 20 in PBS (Millipore Sigma)(PBTw). First, embryos were rehydrated in a solution of 75% methanol to 25% PBTw at room temperature for 5 minutes. This was then followed by the same incubation parameters but 50% methanol to 50% PBTw, then 25% methanol to 75% PBTw. Embryos were then washed in PBTw 3 times for 5 minutes at room temperature and blocked for 30 min using 2% normal goat serum in a solution of 0.1% BSA in PBTw (BBTw). Embryos were incubated with primary antibodies diluted to their appropriate concentrations in BBTw at 4°C for 2 days. Primary was removed and then the embryos washed 5 times for 5 minutes in PBTw at room temperature. Secondary antibody solution was then added to the embryos and incubated for 2 hours at room temperature before being washed 5 times for 5 minutes with PBTw at room temperature. Embryos were then incubated in 500 uL of a 1 ug/mL DAPI solution in PBS for 15 minutes. After DAPI was removed, embryos were kept in PBS at 4°C until mounting. To mount, embryos were transferred to a microscope slide before adding Ultramount Aqueous Permanent Mounting Medium and then coverslipped.

### Western Blots

Adult females were collected and fed for 3-5 days before the ovaries of 30 females were dissected and placed in PBS on ice. The PBS was removed and the ovaries were lysed in 100uL of RIPA lysis buffer (Pierce, ThermoFisher Scientific) with protease inhibitors (cOmplete Mini Protease Inhibitor Cocktail, Roche, Millipore Sigma), PMSF (Roche, Millipore Sigma), and universal nuclease (Pierce Universal Nuclease, Pierce, ThermoFisher Scientific)(83 uL of RIPA, 15 uL of 7x protease inhibitors, 1 uL Unuclease, and 1uL of 100 mM PMSF). The lysate was incubated for 30 min on ice and then centrifuged for 10 minutes at 13000 rpm at 4°C. The supernatant was transferred to a new tube, protein concentration was measured (Pierce BCA Protein Assay Kit, Pierce, ThermoFisher Scientific), and an appropriate volume of supernatant and 2x SDS Protein Gel Loading Solution (Quality Biological) were combined and incubated at 85°C for 2 minutes. Samples were then loaded into a 4-20% Tris-Glycine Gel (ThermoFisher Scientific) and run at 225V for 40 minutes. Proteins were transferred to a 0.2 μm PVDF membrane (ThermoFisher Scientific) at 25V for 2 hours. The membrane was then washed three times in TBS with 0.1% Tween 20 (TBST) and then blocked in TBST containing 5% dried-fat milk (AmericanBio) for 1 hour. The block was removed and membranes were incubated overnight at 4°C in the appropriate primary antibody diluted in TBST containing 5% dried-fat milk (mouse anti-α-Tubulin, 1:10K; mouse anti-HA, 1:1K). Membranes were then washed three times for 5 minutes in TBST and then incubated in secondary antibody diluted in TBST containing 5% dried-fat milk (goat anti-mouse HRP, 1:5K) for one hour.

Secondary antibody solution was then removed and membranes were washed three times for 5 minutes in TBST and then incubated in enhanced chemiluminescence horseradish peroxidase (ThermoFisher Scientific) for 2 minutes before imaging on a Amersham Imager 680 machine (General Electric).

### Image Analysis

Ovaries, testes, and embryos were imaged on a Zeiss 780 LSM confocal microscope (Carl Zeiss AG) using the Zen Black software (Carl Zeiss AG). Image analysis was conducted using Fiji(Schindelin et al. 2012). In order to measure the mean HA staining intensity in *ovo* tagged germariums, single plane 63x germarium images were converted to greyscale for the DAPI and HA channels. A 4.289 area circle in pixels was placed over the center of the nucleus based off of the DAPI staining. The mean staining intensity within this region of interest was then measured for both the DAPI and HA channels. Multiple cells were chosen within each germarium (no less than 2), if that stage was present. For each germarium, two planes of a Z-stack were scored and no less than 60 cells were scored for a given cell stage/genotype. For determining cell stage, cells in the germarium were selected for measurement according to their position in the germarium. Germline stem cells were determined to contain a dot spectrosome indicated by α-Spectrin near the niche of follicle cells at the anterior point of the germarium. Cells in region one contained either a rotated dot spectrosome or clear bar spectrosomes in 2-4 cell cysts and were set in the anterior most portion of the germarium, which lacks follicle cell structure in its outer portion. Cells in region 2a consisted of bar spectrosomes in 8 cell cysts, contained in the area of the germarium where follicle cells begin to form on just the sides of the germarium. Cells in region 2b are contained in a large 16 cell cyst just beginning to be enveloped by follicle cells, and cells in region 3 are in a cyst fully enclosed by follicle cells on the posterior end of the germarium, but have not yet bud off from the rest of the germarium.

### Long and Short Read RNA-seq Analysis

For short read RNA-seq analysis of wild type ovaries, 50 nucleotide single-end reads downloaded from the SRA (SRX2877981-SRX2877983, SRX3291726, SRX2878029-SRX2878031, SRX3291742) (Yang et al. 2018) were mapped to the BDGP Release 6 Drosophila Genome (G. dos Santos et al. 2015) using Hisat2 (-k 1 --rna-strandness R --dta)(Kim et al. 2019). Mapped reads were then sorted and indexed with Samtools (samtools sort and samtools index)(Danecek et al. 2021). In order to generate the read coverage tracks, deepTools’ bamCoverage software (Ramírez et al. 2016) was used to generate a single bigWig file for all wild type ovary replicates (-bs 5 --effectiveGenomeSize 142573017 --normalizeUsing BPM). The bigWig file was then uploaded to UCSC genome browser for visualization(Kent et al. 2002).

PacBio long read mRNA sequencing reads from adult and third instar larval ovaries were downloaded from the SRA (SRX9116734-SRX9116736, SRX9116756, SRX9116757, SRX9116744-SRX9116747) (Brian Oliver 2020). Long reads in fasta format were pooled by tissue type and mapped to the BDGP Release 6 Drosophila genome, with the dm6 NCBI RefSeq GTF file for reference, using SQANTI3 (--skipORF --force_id_ignore)(Tardaguila et al. 2018). Mapped PacBio reads that were PCR duplicates were removed and then filtered by only selecting for reads that mapped upstream of the exon 2 splice acceptor for *ovo* and mapped downstream of the shared stop codon for *ovo*. For filtered reads, the reading frames were then predicted for each transcript based on either the OVO-A or OVO-B start codon, and all reads that were out of frame, or did not utilize the annotated shared stop codon for *ovo*, were removed.

### Statistical Analysis

One-way ANOVA analysis was performed on the mean OVO-HA staining intensities for germ cells within the germarium for the indicated genotypes. R software AOV() function was used to perform and determine ANOVA significance according to the calculated p-value.

## Results

The *ovo* locus encodes a family of C_2_H_2_ zinc-finger transcription factors that result from differential promoter usage. All *ovo* isoforms share a set of four C_2_H_2_ zinc-fingers and stop codons. The somatic version of *ovo*, *svb* is required for viability, and utilizes an upstream promoter that has been shown to be specifically active in somatic cell lineages and not the germline (Mével-Ninio et al. 1995). Two alternative promoters, one utilized by *ovo-A*, and a second utilized by *ovo-B*, have been shown to be specifically active in the germline, and not somatic cell lineages (Figure 1A)(Andrews and Oliver 2002). The germline specific function of *ovo* is believed to be the result of an exon extension in exon 2 (*ovo-RA/RD*), that is not found in *svb*, or annotated to be used by *ovo-A* transcripts (*ovo-RC*, Figure 1A)(Salles et al. 2002; Mével-Ninio et al. 1995). The germline function of this region is supported by the fact that a transposable element insertion in the exon extension found in *ovo^D1rv23^* reverts the strong dominant negative allele, *ovo^D1^*. This is believed to be null for *ovo* as homozygous and hemizygous females show the same phenotype as the antimorph over a deletion/deficiency (Df) (although it should be noted that an antimorph over a Df is also expected to be equivalent to an amorph over a Df). These are *ovo* specific alleles, as hemizygous *ovo^D1rv23^* males are *svb^+^* (B. Oliver, Perrimon, and Mahowald 1987; Salles et al. 2002). We have reexamined the structure/function relationship of the germline *ovo* transcripts, by creating new ‘null’ *ovo* alleles through gene editing, and by studying the locus fine structure with Illumina RNA-Seq (e.g. exon pairs analysis), and with PacBio long-read mRNA-Seq (e.g. “full-length” cDNAs for connectivity) using libraries derived from both adult and third instar larval wild type ovaries.

**Figure 1:**
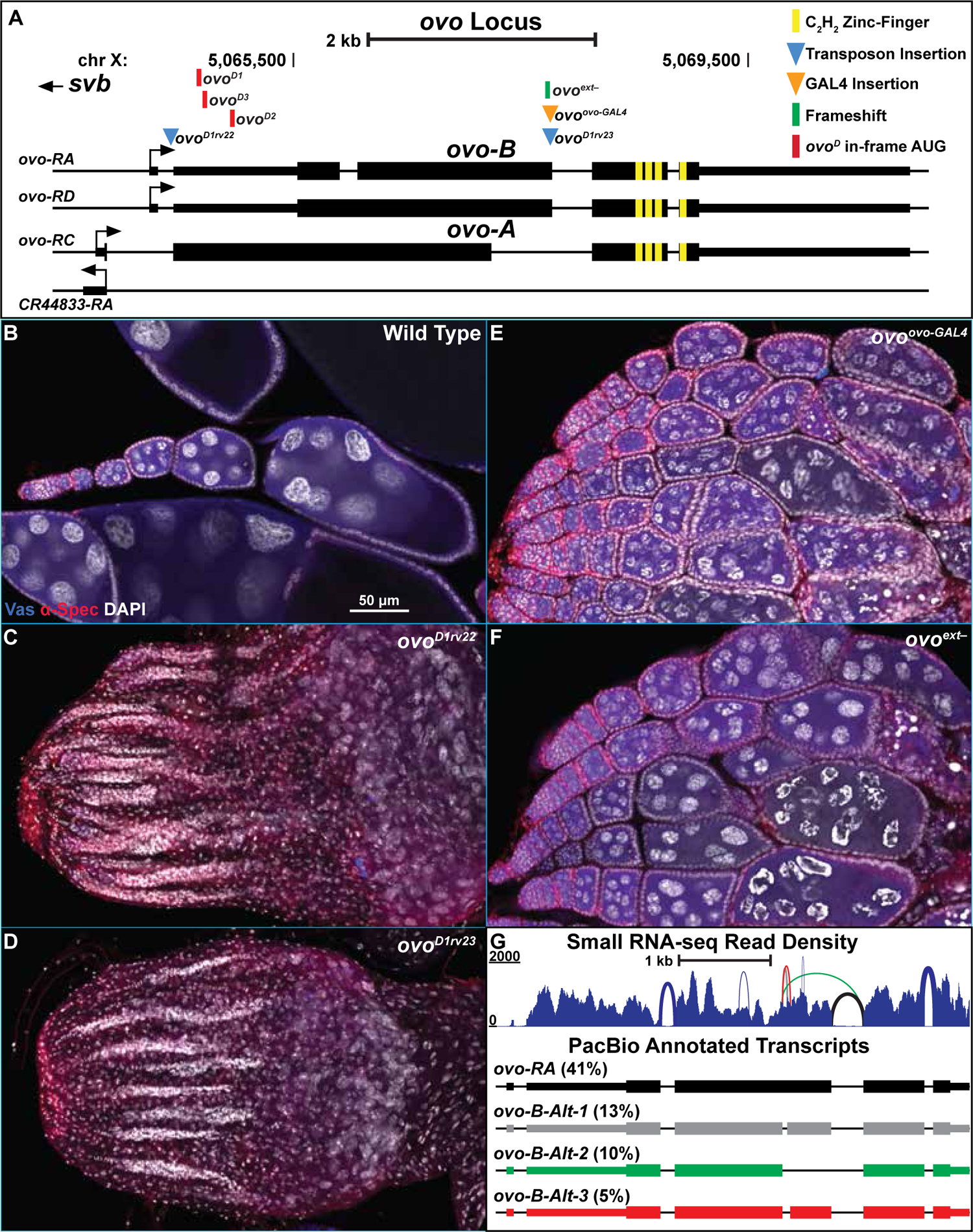
*ovo* locus, phenotypes, and RNA-seq analysis. A) Cartoon of the transcripts expressed from the *ovo* locus according to the current annotations. Green box represents the location of the endogenously generated deletion, orange triangle represents the location of the *T2A-GAL4-3xSTOP* insertion, and blue triangles represent the location of transposon insertions. Yellow boxes represent the location of the shared C_2_H_2_ zinc-fingers and red boxes represent the location of the point mutations for *ovo^D^* alleles. Small rectangles represent untranslated regions, large rectangles represent translated regions. B-F) Immunofluorescent staining of adult ovaries of the indicated genotypes (20x, scale bar = 50 μm). Ovaries were stained for Vas (blue) to label the germline, α-Spectrin (red) to label dot spectrosomes and fusomes, and DAPI to label nuclei. G) Sashimi plot of transcript level small RNA-seq read coverage from wild type ovaries. The weight of the connecting lines across splice junctions correlates with the number of reads that mapped to the splice junction. The most prevalent PacBio transcripts and their percentages among all *ovo-B* transcripts that are supported by small RNA-seq are listed with their annotated name, or novel transcript name.

Revertant alleles of *ovo^D1^* have historically been used to study *ovo* loss-of-function (B. Oliver, Perrimon, and Mahowald 1987; B. Oliver, Pauli, and Mahowald 1990; B. Oliver et al. 1994; Garfinkel, Lohe, and Mahowald 1992; Mével-Ninio et al. 1995; Mével-Ninio, Terracol, and Kafatos 1991; Hinson and Nagoshi 1999; Hinson, Pettus, and Nagoshi 1999; Staab and Steinmann-Zwicky 1996; B. Oliver and Pauli 1998b; Pauli, Oliver, and Mahowald 1993; B. Oliver, Kim, and Baker 1993; Bielinska et al. 2005; Andrews et al. 2000; Lü et al. 1998; Lü and Oliver 2001; Andrews and Oliver 2002). Two alleles specifically, *ovo^D1rv22^* and *ovo^D1rv23^*, are transposable element insertions within the *ovo* locus on the *ovo^D1^*chromosome (Figure 1A)(B. Oliver, Perrimon, and Mahowald 1987; Salles et al. 2002; Mével-Ninio, Mariol, and Gans 1989). *ovo^D1rv22^* is a *Gypsy* insertion between the *ovo-B* specific exon one and two. *ovo^D1rv23^* is a *Stalker2* insertion within the extended exon two of *ovo* (*ovo-RA/RD*). Both alleles are *ovo^-^* and *svb^+^*. The evidence for this is that these alleles are hemi- and homozygous viable, while females are sterile with the same phenotype as *ovo^D1^/Df* (B. Oliver, Perrimon, and Mahowald 1987). Phenotypically, both alleles behave the same. These alleles have been well characterized, and our work confirmed that *ovo^D1rv22^* and *ovo^D1rv23^* ovaries are 100% agametic (50/50), only containing strings of soma-only ovarioles as seen with DAPI staining (Figure 1C,D, Table S2). Based on this evidence, both alleles have been considered to be amorphic *ovo* alleles. However, given that *ovo^D1^/Df* and *ovo^-^/Df* have the same phenotype, it is impossible to determine if they are truly null or weak (recessive) antimorphs.

### Does *ovo* germline function require both extended exon and non-extended exon isoforms?

In our ongoing efforts to revisit *ovo*, we decided to generate new endogenous alleles that would specifically disrupt *ovo* and not *svb* function, without potentially confounding effects of *ovo^D1^*. Simultaneously, we also wanted the ability to express *GAL4* in the same expression pattern as *ovo* while also disrupting wild type *ovo* function. To do this, we placed a *T2A-GAL4-3xSTOP* DNA sequence (Daniels et al. 2014; Brand and Perrimon 1993) one codon upstream of the splice site in the extended exon two of the annotated *ovo-B* transcripts (Figure 1A) with CRISPR/Cas9. In this design, translation begins at the endogenous *ovo-B* start codon, and the ribosome skips the peptide bond once it reaches the viral *T2A* sequence (Daniels et al. 2014), before continuing to translate the downstream GAL4 reading frame and eventually cease translation at the downstream *3xSTOP* codons in the *GAL4* sequence. This would have the functional consequence of producing a DNA-binding domain lacking N-terminal OVO-B and a separate GAL4 protein. The downstream shared zinc-fingers of OVO would not be translated. If *ovo^D1rv23^*is a null, then this new allele should result in the same phenotype.

After endogenously inserting the *T2A-GAL4-3xSTOP* with CRISPR/Cas9, resulting in the *ovo^ovo-GAL4^* allele (Figure 1A), we found that males and females hemi- and homozygous for the insertion were viable and showed no overt somatic phenotypes, indicating that the *T2A-GAL4-3xSTOP* insertion was not disrupting *svb* transcripts, as expected. We also found that homozygous females were sterile, indicating that the exon two extension is required for *ovo* function. However, the *ovo^ovo-GAL4^*phenotype was much weaker than we expected. Based on the ovarian phenotype of *ovo^D1rv23^* ovaries, we expected *ovo^ovo-GAL4^* ovaries would also be agametic. However, *ovo^ovo-GAL4^* ovaries always (50/50) contained germ cells (Figure 1E, Table S2). *ovo^ovo-GAL4^*ovaries contained strings of germ cell containing ovarioles with wild type appearing germariums and developing egg chambers that arrested around stage 5 of oogenesis, based on the ‘blob’ chromosome morphology seen in stage 5 egg chambers (Dej and Spradling 1999). There were no later egg chamber stages, suggesting developmental arrest or death at stage 5. We observed evidence of degenerating egg chambers based on the presence of pyknotic nuclei suggesting that stage 5 *ovo^ovo-GAL4^*egg chambers die rather than accumulate. This unexpectedly weak *ovo* phenotype brought into question our current view of the *ovo* null phenotype and thus the function of *ovo* in the germline.

Since the insertions in *ovo^ovo-GAL4^* and *ovo^D1rv23^*are disrupting the same annotated *ovo-B* transcripts, but resulted in dramatically different female germ cell phenotypes, there were three plausible explanations. One, *ovo* is not required for female germline viability; two, there are *ovo* transcripts that do not have the exon extension; and three, there is an unexpected effect of GAL4 leading to germ cell survival. In order to address this last (unlikely) scenario first, we designed two gRNAs targeting the exon extension and used CRISPR/Cas9 to create mutations specifically in the exon extension of *ovo*. We recovered an X chromosome containing a single nucleotide deletion in the fourth to last codon in the extended exon 2 before the splice site, that would result in a frameshift and early stop codon 15 codons downstream. This new stop codon would be upstream of the four shared C_2_H_2_ zinc-fingers of *ovo*/*svb*, however, only isoforms containing the exon extension would be affected. This allele we called *ovo^ext–^* (Figure 1A). If the weak germline phenotype was due to ectopic expression of GAL4 in *ovo^ovo-GAL4^*, then *ovo^ext–^* would not phenocopy it, and would have the germ cell death phenotype observed in *ovo^D1rv23^* ovaries.

Immunostaining of *ovo^ext–^* ovaries showed that germ cells did survive, similar to *ovo^ovo-GAL4^* ovaries (Figure 1F, Table S2). However, the presence of egg chambers differentiating slightly past stage 5 were observed, based on chromosome morphology visualized with DAPI. These egg chambers did eventually degenerate, seen with the presence of abnormal chromosome morphology as well as pyknotic nuclei in the germline when compared to wild type controls. This result confirmed that the *ovo^ovo-GAL4^*germ cell phenotype was not due to the presence of ectopically expressed GAL4. Thus, both of these alleles fail to phenocopy *ovo^D1rv23^*, even though they all disrupt the same *ovo-B* isoforms. The true nature as to why *ovo^D1rv23^* ovaries do not contain germ cells was still a mystery.

### Reannotation of the *ovo* locus

Previous work has shown that *ovo-B* transcripts utilizing both the exon extended splice variant as well as the non-extended exon two splice variant, similar to *svb*, are present in germ cells (Salles et al. 2002).

Whereas *svb* transcripts have only been shown to utilize the non-extended exon two isoform. Furthermore, one copy of an *ovo* genetic construct containing only the non-extended exon isoform could rescue germ cell death, and allow for egg chamber development up to stage 4 in “*ovo^-^*“, *ovo^D1rv^* ovaries. Two copies of the same genetic construct allowed for similar rescue with an increase in later stage egg chamber differentiation (Salles et al. 2002). These data support the idea that *ovo-B* might be utilizing the non-extended exon splicing variant that is similar to *svb/ovo-A*, since the germ cell survival but arrested egg chamber phenotype in *ovo^ovo-GAL4^* and *ovo^ext–^* ovaries is similar to containing only the non-extended exon *ovo* rescue genetic construct in *ovo^D1rv^* ovaries. If this is true, then the current FlyBase annotation of the *ovo* locus is incorrect.

If *ovo-B* is utilizing other splicing variations than what is currently annotated, the evidence for this should be present in datasets from modern RNA sequencing (RNA-seq) methods. To address this transcript variation at the *ovo* locus, we analyzed small RNA-seq (Yang et al. 2018) and long-read RNA-seq (Brian Oliver 2020) from wild type *Drosophila* ovaries from the SRA (Table S1). We mapped PacBio long-read mRNA transcripts derived from both adult and third instar larval wild type ovaries and filtered for full length transcripts belonging to the *ovo* locus (see methods). These were further filtered to remove transcripts that resulted in out-of-frame zinc-fingers and thus left us with a total of 5,079 in-frame, potentially full-length transcripts from the *ovo* locus (Table S3) to use for comprehensive reannotation of the *ovo* locus.

Since we filtered for cDNAs starting upstream of the shared exon 2 splice acceptor site, we could potentially capture any variations in splicing in either *ovo-A* or *ovo-B*. The *ovo-B* transcripts are currently annotated to contain a small intron (chrX: 5,065,913-5,066,073) that corresponds to FlyBase annotation isoform *ovo-RA*, or are annotated to not contain this intron within exon 2, which corresponds to isoform *ovo-RD* (Figure 1A)(Gramates et al. 2022; G. dos Santos et al. 2015). We checked the abundance of this splice junction usage in the PacBio long-read transcripts and found that 94% (4,913/5,079) of transcripts contained the annotated intron in exon 2 that corresponds to isoform *ovo-RA*, and thus this is the predominant splicing pattern. Furthermore, we found that the splicing of the annotated exon 3 to exon 4 splice and the annotated exon 4 to exon 5 splice in *ovo-RA*, was used 86% (4,371/5,079) and 99% (5,070/5,079) of the time, respectively. This indicates that there was little variation in splicing of the zinc-finger containing exons 4 and 5. However, there seemed to be a greater variety of splicing patterns in the annotated exon 3 of *ovo-RA*.

Looking at all transcripts derived from the *ovo-B* promoter (4,913 in total), we found that the most prevalent transcript in the PacBio long-read RNA-seq data was the annotated *ovo-B* transcript (*ovo-RA*) which accounted for 41% (2,002/4,913) of all *ovo-B* transcripts (Figure 1G, *ovo-RA*). The next most prevalent isoform contained a small intron in exon 3 (chrX:5067247-5067302, Figure 1G, *ovo-B-Alt-1*) which accounted for 13% (632/4,913) of all *ovo-B* transcripts. The third most prevalent transcript that accounted for 10% (489/4,913) of the *ovo-B* reads used the *ovo-RC* splice site, which is annotated to be specific to *ovo-A* and *svb*, but not utilized by *ovo-B* (Figure 1G, *ovo-B-Alt-2*). It is this *ovo-B-Alt-2* isoform that is important for understanding the phenotype of the exon two extension alleles. Assuming that the mutations have no effect on transcription of *ovo* (which is unlikely given the autoregulatory function of OVO), there are wild type OVO transcription factors produced in *ovo^ovo-GAL4^*and *ovo^ext–^* ovaries, and antimorphic OVO^D1^ transcription factors produced in “null” *ovo^D1rv23^* ovaries.

In order to examine the fine structure of the locus with another method, we asked if there was support for the splicing variations found in the long-read dataset, in small RNA-seq reads derived from *Oregon-R* and *w^1118^* adult ovaries (Yang et al. 2018). After mapping small RNA-seq reads, we did not have evidence for any reads that mapped to the *ovo-A* promoter or *ovo-A* exon 1 to exon 2 splice junction (consistent with them being rare, which we also observed in the full length cDNAs), however we did have evidence for reads mapping to the *ovo-B* promoter. Therefore, we believed that a vast majority of reads mapping to *ovo* transcripts are likely a result of *ovo-B* specific transcripts. We found that small RNA-seq reads supported the predominant annotated splicing pattern from exon 3 to 4, corresponding to the *ovo-RA* transcript. Small RNA-seq reads also supported a third PacBio transcript that was found in 5% (247/4,913) of all *ovo-B* specific PacBio reads (Figure 1G, *ovo-B-Alt-3*). Other splicing variations were found within the small RNA-seq read dataset, however, none of these other splicing variations were found to be in the top ten most abundant PacBio reads for *ovo-B*, and although they might exist *in vivo*, they are likely to be rare. Most importantly, small RNA-seq reads supported the second most abundant PacBio transcript small intron variant (Figure 1G, *ovo-B-Alt-1*) as well as the *ovo-RC*-like *ovo-B* splicing variant that excludes the extended exon from exon 3 to 4 (Figure 1G, *ovo-B-Alt-2*). This last result confirms that *ovo^ovo-GAL4^* and *ovo^ext–^* are not molecular nulls. They encode minor *ovo-B^+^* transcripts.

The structure and function of *ovo-A* has been defined almost entirely by the activity of the dominant antimorphs (*ovo^D1^, ovo^D2^, ovo^D3^, ovo^D4^,* and *ovo^D5^*) which append OVO-A codons to the N-terminus of OVO-B as a result of new initiation codons (Mével-Ninio et al. 1996; Andrews, Levenson, and Oliver 1998). We still do not know how abundant OVO-A might be and what it does in wild type flies. Of the 5,079 full length transcripts that mapped to *ovo*, 49 (1%) mapped specifically to the *ovo-A* promoter. These *ovo-A* transcripts showed similar splicing patterns to *ovo-B*, with the use of the small intron in exon 2 to be found in 86% (42/49) of transcripts. We found that the most abundant *ovo-A* isoform, accounting for 47% (23/49) of transcripts, utilized the same splicing pattern within the open reading frame as *ovo-RA*. The rest of the *ovo-A* specific isoforms utilized a variation of splicing patterns in exon 3, which we also found for *ovo-B* transcripts. Additionally, all *ovo-A* transcripts in the PacBio dataset faithfully used the same extended exon 3 into exon 4 splice junction as *ovo-RA*. There was not any evidence for the annotated *ovo-RC* isoform in this dataset. Therefore, *ovo-A* transcripts using the same exon 3 splice pattern as *svb*, are incredibly rare, if they exist at all.

To briefly summarize, our analysis integrating the genetics of *ovo^ovo-GAL4^* and *ovo^ext–^* alleles, along with the RNA-seq transcripts expressed from *ovo*, suggested that *ovo* utilizes both the extended exon splice site as well as the non-extended exon splice site (Figure 2A). This suggests that *ovo^D1rv23^* is a recessive antimorph expressing low levels of OVO^D1^. Our newly generated alleles *ovo^ovo-GAL4^*and *ovo^ext–^* are likely hypomorphs.

**Figure 2:**
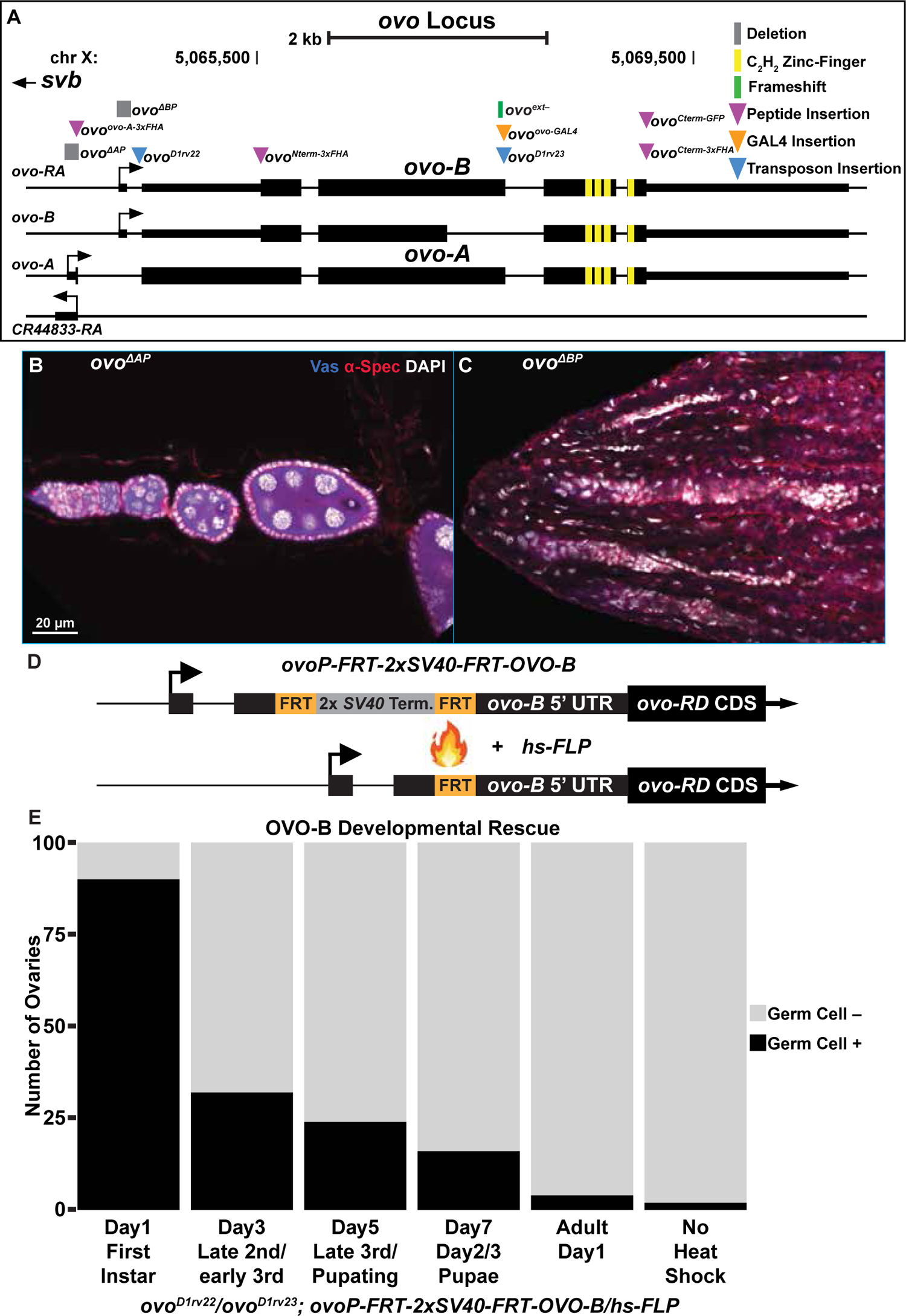
Loss of Function OVO-A and OVO-B in the Adult and Developing Germline. A) Cartoon of the transcripts expressed from the *ovo* locus updated based on the genetic and RNA-seq results. Green boxes represent the location of endogenously generated deletions, orange triangle represents the location of the *T2A-GAL4-3xSTOP* insertion, blue triangles represent the location of transposon insertions, and purple triangles represent the location of small peptide insertions. Yellow boxes represent the location of the shared C_2_H_2_ zinc-fingers and red boxes represent the location of the point mutations for *ovo^D^*alleles. Small rectangles represent untranslated regions, large rectangles represent translated regions.B-C) Immunofluorescent staining of adult ovaries of the indicated genotypes (40x, scale bar = 20 μm). Germariums were stained for Vas (blue) to label the germline, and α-Spectrin (red) to label dot spectrosomes and fusomes, and DAPI (white) to label nuclei. D) Cartoon of *ovoP-FRT-2xSV40-FRT-OVO-B* transgenic OVO rescue construct. In the presence of *FLP*, such as under the control of a heat shock promoter (*hs-FLP*), recombination will occur between the *FRT* sites and thus remove the 2x*SV40* termination sequences, allowing for expression of the downstream OVO-B reading frame. Small rectangles represent untranslated regions, large rectangles represent translated regions. E) OVO-B developmental rescue timeline. Flies with the genotype *ovo^D1rv22^*/*ovo^D1rv23^*; *ovoP-FRT-2xSV40-FRT-OVO-B*/*hs-FLP* were heat shocked at the indicated time point during development. Flies were allowed to continue to develop and adults were scored for the presence of germ cell containing ovarioles or ovarioles not containing germ cells.

### *ovo-A* and *ovo-B* null allele

The *ovo^D1rv23^* allele shows the same “*ovo* null” or recessive antimorphic adult female germ cell death phenotype as *ovo^D1rv22^*. While this allele shares the *ovo^D1^* background with *ovo^D1rv22^*, *ovo^D1rv22^* results from a *Gypsy* insertion near the *ovo-B* promoter. The *ovo* locus is a hotspot for *Gypsy* insertions which disrupt the germline function of *ovo* and result in an orientation-dependent lozenge-like rough-eye phenotype (Mével-Ninio, Mariol, and Gans 1989; Mével-Ninio, Terracol, and Kafatos 1991; Dej et al. 1998). *Gypsy* retrotransposons contain insulator DNA sequences that have been shown to negatively influence gene expression by disrupting necessary enhancer-promoter interactions (Gdula, Gerasimova, and Corces 1996). It is possible that the *Gypsy* insertion in *ovo^D1rv22^* is disrupting the expression of all *ovo* transcripts through this type of mechanism. Given this long list of potential confounding factors, it is not at all clear if *ovo^D1rv22^* is null or a recessive antimorph. In order to address this, we created *ovo-A* and *ovo-B* promoter deletions using updated genetic techniques to better understand the functions of these two classes of germline restricted transcripts.

Our understanding of *ovo* promoters stems from transgene studies (B. Oliver et al. 1994; Mével-Ninio et al. 1995; Hinson, Pettus, and Nagoshi 1999; Bielinska et al. 2005; Salles et al. 2002; Lü et al. 1998; Lü and Oliver 2001; Andrews et al. 2000; Andrews and Oliver 2002). These reporters indicate that *ovo* is specifically expressed in the *Drosophila* germline and they respond to *ovo* copy number *in trans*. Deletion of the *ovo-A* promoter in a genetic rescue construct complements the germ cell death phenotype of *ovo^D1rv^* germ cells (Andrews et al. 2000), indicating that OVO-B is sufficient for the rescue. However, females expressing no *ovo-A* showed a grandchildless phenotype, suggesting a maternal role in germline development. As with most phenotypes, this grandchildless phenotype was in the background of a weak dominant-negative *ovo^D1^*. Thus, that maternal phenotype could easily have been due to a maternal effect of too much OVO-A-like, OVO^D1^. In contrast to *ovo-A* transgenes, deleting the *ovo-B* promoter in a reporter construct showed a great loss of reporter expression, while this same deletion in a genetic rescue construct was unable to rescue the cell death of *ovo^D1rv^* germ cells (Andrews and Oliver 2002).

We wanted to repeat these experiments in the context of endogenous *ovo^+^* to determine the phenotypes of female germ cells that lack either *ovo-A* or *ovo-B*. Therefore, we designed CRISPR/Cas9 gRNA and homology-directed repair constructs to delete both the *ovo-A* and *ovo-B* promoters, resulting in new alleles called *ovo^ΔAP^* and *ovo^ΔBP^*, respectively (Figure 2A).

When we deleted the endogenous *ovo-A* promoter (*ovo^ΔAP^*), we found that males and females were both viable and fertile. This result was consistent with no role for the *ovo-A* promoter in *svb* or *ovo-B* function. We immunostained homozygous *ovo^ΔAP^* ovaries and found that germarium morphology throughout egg chamber development looked wild type (Figure 2B) in 100% of *ovo^ΔAP^* ovaries scored (100/100, Table S4). Of course, there could be defects that we failed to observe, but there was no overt function of OVO-A-Previous work on *ovo* has shown that *ovo-B* rescue transgenes with a deletion of the *ovo-A* promoter in an *ovo^D1rv^* background showed a maternal grandchildless phenotype with between 10-40% penetrance, which underwent gradual extinction (Andrews and Oliver 2002). We tested whether *ovo^ΔAP^* mothers crossed to wild type fathers resulted in a grandchildless phenotype. We found that 98% (98/100, Table S4) of ovaries in *ovo^ΔAP^*/+ daughters contained germ cells and therefore does not suggest a pervasive maternal grandchildless phenotype of *ovo-A^-^.* The OVO-A transcription factor is a powerful repressor, so it is possible that the OVO-A encoding OVO^D1^ in *ovo^D1rv^* females has a maternal effect. Indeed, this might explain the very early germline defect in one of the early reports on an *ovo* requirement in the embryo (B. Oliver, Perrimon, and Mahowald 1987), although this early phenotype has never been replicated despite efforts to do so (Rodesch et al. 1995). Obviously, exploring the maternal effect of a gene absolutely required for egg development poses technical challenges that will require elegant genetic engineering to address. We have no evidence here that removing OVO-A had any negative effect on female germ cells or the progeny from those females.

In stark contrast to the effect of deletion of the *ovo-A* promoter, deletion of the *ovo-B* promoter (*ovo^ΔBP^*), had a drastic effect on ovarian morphology. Hemi and homozygous *ovo^ΔBP^* flies were viable, indicating that the deletion did not disrupt *svb* function, and males were fertile, consistent with previous work on *ovo^D1rv^* (B. Oliver, Perrimon, and Mahowald 1987). However, homozygous *ovo^ΔBP^* females were completely sterile. Immunostaining of *ovo^ΔBP^*ovaries showed a complete loss of the germline in 100% of females tested (100/100, Table S4), indicated by the lack of Vas positive cells, with only remnants of the somatic ovariole structure seen by DAPI positive cells localized anteriorly to posteriorly (Figure 2C). This finding indicates that *ovo-B* is required for female germ cell viability, as has been reported previously in the *ovo^D1rv^*alleles (Salles et al. 2002; Andrews and Oliver 2002). Importantly, the homozygous and hemizygous *ovo^ΔBP^* phenotype is the same as seen in *ovo^D1^/Df*. This indicates that the *ovo^-^* phenotype is indeed female germline death.

Work on both the *ovo^D1^ and ovo^D1rv^* alleles indicate that the dominant-negative functions can disrupt female germline development throughout the lifecycle. It does not necessarily follow that OVO-B is required throughout the lifecycle. Deletion of the *ovo-B* promoter resulted in a pervasive female specific germ cell death phenotype in adults. We were interested in determining the developmental stage(s) where OVO-B is required in the female germline. While *ovo^D1^/+* are sterile, mitotic +/+ clones in those females are capable of forming fertile eggs. The earlier those clones are induced, the higher the recovery of wildtype germline clones (Perrimon 1984). This is consistent with an ongoing requirement for *ovo*.

Previous work has shown a reduction in germ cell number in *ovo^D1rv^*females during the larval and pupal stages (Staab and Steinmann-Zwicky 1996). In order to more directly determine the requirement for OVO-B during development, we created a temporally activated *ovo^+^* rescue transgene.

The *ovo-B* rescue transgene consisted of the upstream regulatory enhancer elements that are sufficient for *ovo* expression in the female germline (B. Oliver et al. 1994; Mével-Ninio et al. 1995): the *ovo-B* promoter, the *ovo-RD* cDNA, and the *ovo* 3’ UTR. However, a *FRT*-2x*SV40* terminator-*FRT* DNA sequence was placed within the untranslated exon 2 region of *ovo*, resulting in *ovoP-FRT-2xSV40-FRT-OVO-B* (Figure 2D). Under normal circumstances, transcription of the *ovo* rescue transgene should terminate when the RNA polymerase reaches the tandem *SV40* termination sequence (Grass, Jove, and Manley 1987), resulting in early transcriptional termination of the transgene. However, when FLP recombinase is expressed, such as under the control of a heat shock promoter (*hs-FLP*), then the FLP recombinase would catalyze a recombination event between the two *FRT* sites (Golic and Lindquist 1989), and thus remove the tandem *SV40* termination sequences, allowing full transcription of the transgene (Figure 2D). We used the *ovo^D1rv^* background to reduce any potential leaky expression of *ovo-A.* To determine if this transgene behaved as intended, we scored the number of germ cell containing ovaries in *ovo^D1rv22^*/*ovo^D1rv23^*; *ovoP-FRT-2xSV40-FRT-OVO-B*/*hs-FLP* females. We found that 2% of females had germ cell containing ovaries (2/100, Figure 2E), thus indicating that there was little to no expression of the downstream OVO-B cDNA when the tandem 2x*SV40* termination sequence remained in place.

To explore the temporal requirement, we heat shocked either one day after egg laying, and in two day developmental increments after day one until adulthood. We found that 90% (90/100) of female adult flies that were heat shocked one day after egg laying had ovaries containing germ cells (Figure 2E). This indicated that removing the tandem *SV40* termination sequence through FLP-mediated recombination in our *ovo^+^* rescue transgene allowed for sufficient *ovo-B* expression and OVO^+^ activity to rescue the *ovo^-^* germ cell phenotype in some females, but in the minority, the rescue was too late. The percent of adult females that had germ cell containing ovaries when we expressed *ovo-B* at 3, 5, and 7 days post-egg laying, as well as day-one adults, were 32%, 24%, 16%, and 4%, respectively (Figure 2E). This gradual decrease in rescue suggests that germ cells gradually succumb to the absence of OVO-B, but once *ovo-B* expression is restored, enough germ cells recover and are able to proliferate to populate an adult ovary. This polyphasic requirement for OVO raises the hypothesis that there is a maternal contribution. The gradual decrease in rescue could be due to perdurance of maternally deposited OVO for example.

### OVO-B extended and short exon isoforms have distinct functions

Now that we had bonafide *ovo-B^-^* alleles, we could re-examine the exon extension alleles to explore their genetic function. The locus re-annotation and weak *ovo* phenotypes suggest that *ovo^ovo-GAL4^*and *ovo^ext–^* are partial loss-of-function alleles. Our *ovo* small and long-read RNA-seq transcript analysis confirmed that there are alternatively spliced *ovo-B* isoforms that do not contain the extended exon, thus allowing for the possibility that the presence of these non-extended exon OVO protein isoforms contain sufficient OVO^+^ activity for germ cell survival. However, it is unclear whether the germ cell survival, but arrested egg chamber phenotype is due to lower levels of OVO, or if there is a specific requirement for long and short OVO-B isoforms. We know that expressing the non-extended exon isoform rescues *ovo^D1rv^*germ cell death, but not full fertility. Extra copies of the rescue construct *in trans* resulted in further rescue of the *ovo^D1rv^* phenotype compared to just one copy (Salles et al. 2002). A similar genetic rescue construct that contained both the extended and non-extended exon isoforms was able to fully rescue *ovo^D1rv^* germ cells to wild type. Since extra copies of the non-extended exon isoform *in trans* increased the phenotypic rescue, it is possible that the absolute level of OVO-B, and not the alternative OVO-B protein isoforms is sufficient to rescue the *ovo-B^-^*germ cells.

Since the insertion of *T2A-GAL4-3xSTOP* in *ovo^ovo-GAL4^*specifically disrupts the extended exon *ovo* isoforms, and not the non-extended *ovo* isoforms, we asked if it behaved as a partial loss-of-function allele over *ovo-B^-^. ovo^ovo-GAL4^* ovaries contained germ cells in 100% of ovaries scored, however, if *ovo^ovo-GAL4^* was transheterozygous with *ovo^ΔBP^*, we found that only 52% of ovaries contained Vas positive germ cells with egg chambers arresting at stage 5. This indicates that *ovo* activity is low in *ovo^ovo-GAL4^.* If this is correct, then we could use the *GAL4* expressed under *ovo* control to test rescuing cDNAs. This is an ideal background for testing both the positive and negative effects of various cDNAs. We drove *UASp-GFP* (control), *UASp-3xFHA-OVO-A* (expressing the repressive OVO isoform), *UASp-3xFHA-OVO-B^ext+^*(expressing the extended exon OVO-B isoform), or *UASp-3xFHA-OVO-B^ext–^*(expressing the non-extended exon OVO-B isoform) and examined the ovarian phenotype by counting the number of Vas^+^ stage 5 egg chamber containing ovarioles per ovary (Figure 3A).

**Figure 3:**
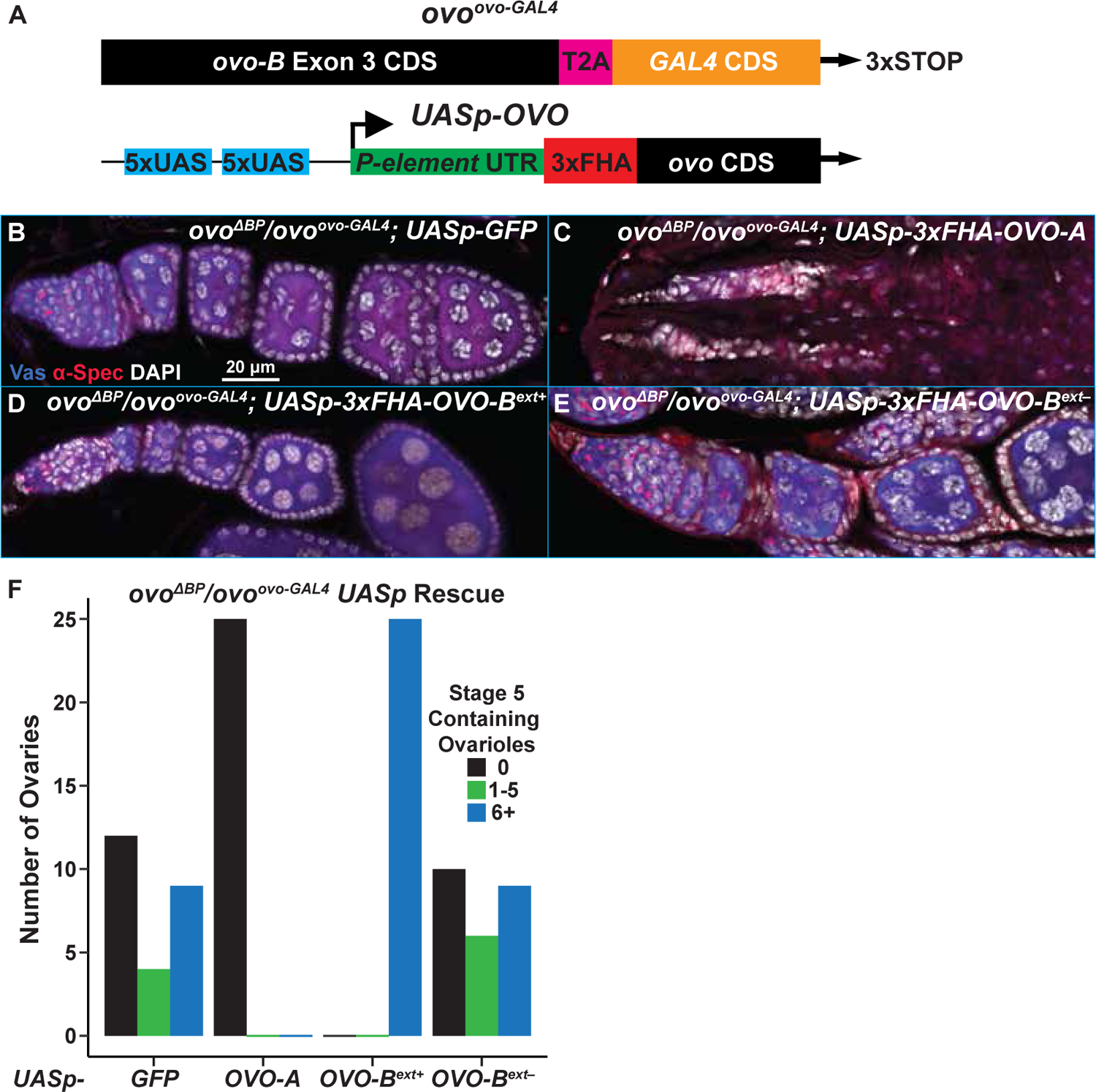
OVO Isoform rescue of *ovo^ovo-GAL4^*/ovo*^ΔBP^* Ovaries. A) Cartoon of *ovo^ovo-GAL4^* allele and *UASp* rescue transgenes. Small rectangles represent untranslated regions, large rectangles represent translated regions. B-E) Immunofluorescent staining of adult ovaries of the indicated genotypes (40x, scale bar = 20 μm). B-E) Ovarioles were stained for Vas (blue) to label the germline, α-Spectrin (red) to label dot spectrosomes and fusomes, and DAPI (white) to label the nuclei. F) Graph of the number of ovo*^ΔBP^*/ovo*^ovo-GAL4^* ovaries that contained Vas positive germarium through stage 5 egg chambers paired with the respective *UASp* transgene.

Immunostaining of the control *ovo^ovo-GAL4^*/*ovo^ΔBP^*; *UASp-GFP* ovaries showed that surviving germ cells did form ovariole structures containing egg chambers which arrested at stage 5 based on the characteristic ‘blob’ chromosome structure of nurse cells that was visualized through DAPI staining (Figure 3B). We found that 12/25 ovaries did not contain Vas positive germ cells, 4/25 contained 1-5 ovarioles with Vas positive germ cells, and 9/25 ovarioles contained six or more ovarioles with Vas positive germ cells (Figure 3F). We expected that ectopic OVO-A expression in *ovo^ovo-GAL4^*/*ovo^ΔBP^*ovaries would exacerbate the phenotype, as ectopic expression of OVO-A causes germ cell defects, presumably by negative autoregulation of *ovo-B* (Andrews and Oliver 2002). Similarly, the *ovo^D^* class of mutations are dominant female sterile, due to the addition of the repressive OVO-A transcriptional domains onto OVO-B proteins, and result in negative autoregulation of *ovo (Andrews, Levenson, and Oliver 1998; Mével-Ninio et al. 1996)*. Immunostaining of ovo*^ΔBP^*/ovo*^ovo-GAL4^; UASp-3xFHA-OVO-A* ovaries never contained stage 5 egg chambers, and never more than six Vas^+^ containing ovarioles (Figure 3C, 3F), like ovo*^ΔBP^* ovaries. There were strings of DAPI positive somatic cells localized anteriorly to posteriorly that lacked any evidence of germ cells. Surprisingly, 52% of ovo*^ΔBP^*/ovo*^ovo-GAL4^; UASp-3xFHA-OVO-A* ovaries contained at least one Vas positive germ cell. Thus, while *ovo-A* exacerbated the ovo*^ΔBP^*/ovo*^ovo-GAL4^*ovary phenotype, that phenotype was slightly weaker than seen in *ovo^ΔBP^*ovaries.

Next, we addressed the ability of different *ovo-B* isoforms to rescue ovo*^ΔBP^*/ovo*^ovo-GAL4^* ovaries. Immunostaining ovo*^ΔBP^*/ovo*^ovo-GAL4^*; *UASp-3xFHA-OVO-B^+^*resulted in a full rescue of the arrested egg chamber phenotype in ovo*^ΔBP^*/ovo*^ovo-GAL4^*female ovaries (Figure 3D, 3F). The ovaries of these females contained egg chambers of all differentiation stages based on the nuclear morphology seen with DAPI. Additionally, females of this genotype were able to lay eggs. This result indicated that expression of OVO-B containing the extended exon downstream of GAL4 expressed from the *ovo* locus was able to fully rescue. However, female ovo*^ΔBP^*/ovo*^ovo-GAL4^*; *UASp-3xFHA-OVO-B^ext–^* flies that only expressed the non-extended exon OVO-B isoform, were not rescued. These ovaries were indistinguishable from the ovo*^ΔBP^*/ovo*^ovo-GAL4^*; *UASp-GFP* control ovaries (Figure 3E). Specifically, many ovo*^ΔBP^*/ovo*^ovo-GAL4^*; *UASp-3xFHA-OVO-B^ext–^*ovaries did not contain Vas positive germ cells (10/25), 6/25 contained 1-5 ovarioles with germ cells, and only 9/25 ovarioles contained six or more ovarioles with germ cells (Figure 3F). This suggested that the OVO-B extended exon isoforms, encoding a longer OVO-B isoform, are a qualitative requirement for oogenesis, not a quantitative requirement. We conclude that long and short OVO-B isoforms have distinct functions.

### OVO is expressed in the female and male germline

Given that the *ovo* locus is required in multiple stages of germline development, we hypothesized that OVO-B was expressed throughout female germline development. Unfortunately, we do not have OVO antibodies or tagged endogenous OVO-A or OVO-B. To bypass this, we used CRISPR/Cas9 to place an N-terminal 3xFlag-3xHA (3xFHA) tag immediately downstream of the start codon of OVO-A (*ovo^ovo-A-3xFHA^*). This tag were also placed immediately downstream of the start codon of OVO-B (*ovo^Nterm-3xFHA^*) as well as immediately upstream of the stop codon in both the OVO-A and OVO-B reading frames (*ovo^Cterm-3xFHA^*, *ovo^Cterm-GFP^*). Therefore, the OVO-A N-terminal tag should be specific to only OVO-A protein, while the tags placed downstream of the OVO-B start codon and upstream of the shared stop codon, would tag OVO-A internally and OVO-B at the N-terminus, or both isoforms at the C-terminus. None of these tags are OVO-B specific. Although it has been shown that OVO-B is the predominant protein isoform expressed in the female germline (Hayashi et al. 2017; Andrews and Oliver 2002), we wanted to ensure that we could detect only OVO-B and therefore used CRISPR/Cas9 to delete the *ovo-A* promoter in both the _*ovo*_*_Nterm-3xFHA_* and _*ovo*_*_Cterm-3xFHA_* alleles. This resulted in _*ovo*_*_ΔAP-Nterm-3xFHA_* and _*ovo*_*_ΔAP-Cterm-3xFHA_* alleles, which should only tag the germline OVO-B, but not OVO-A, protein isoforms (in addition to SVB in the soma).

All these alleles were viable and fertile, indicating that *ovo*/*svb* function was not perturbed and thus these alleles could be used to study the expression and localization patterns of endogenous OVO. Our endogenous OVO tagged isoforms (minus *ovo^ovo-A-3xFHA^*) will also inherently tag SVB, so we cannot rule out that detection of our tags is purely due to OVO and not SVB. However, *ovo* and *svb* are genetically separable. Alleles that specifically disrupt the *svb* promoter/exon are phenotypically *svb^-^ ovo^+^*, and alleles specifically disrupting the *ovo* promoter display a *svb^+^ ovo^-^* phenotype (B. Oliver, Perrimon, and Mahowald 1987). Furthermore, reporter constructs containing only the regulatory regions and promoters for *ovo*, and not *svb*, are strictly expressed in the germline (Salles et al. 2002; Andrews and Oliver 2002). Therefore, *svb* and *ovo* use distinct regulatory sequences to ensure that their expression is cell type specific, and detection of tags in the germline will likely be specific to OVO.

Since OVO is required for female fertility, we decided to look at localization in adult ovaries first. We _immunostained homozygous *ovo*_*_Nterm-3xFHA_*_, *ovo*_*_Cterm-3xFHA_*_, *ovo*_*_ΔAP-Nterm-3xFHA_*_, and *ovo*_*_ovo-A-3xFHA_* _ovaries_ (Figure 4A-D) for anti-HA immunoreactivity and found that in all four genotypes, HA staining was restricted specifically to the germline within the ovaries due to its expression in only Vas positive cells. Also, in all genotypes, localization was found to be enriched in the nucleus versus the cytoplasm, since it overlapped with the DAPI DNA counterstain, as opposed to overlapping with perinuclear/cytoplasmic Vas. These results are consistent with OVO transcription factor function in the germline (B. Oliver, Perrimon, and Mahowald 1987; Andrews and Oliver 2002), and sequence-specific DNA binding *in-vitro* (Lee and Garfinkel 2000; Bielinska et al. 2005; Lü and Oliver 2001).

**Figure 4:**
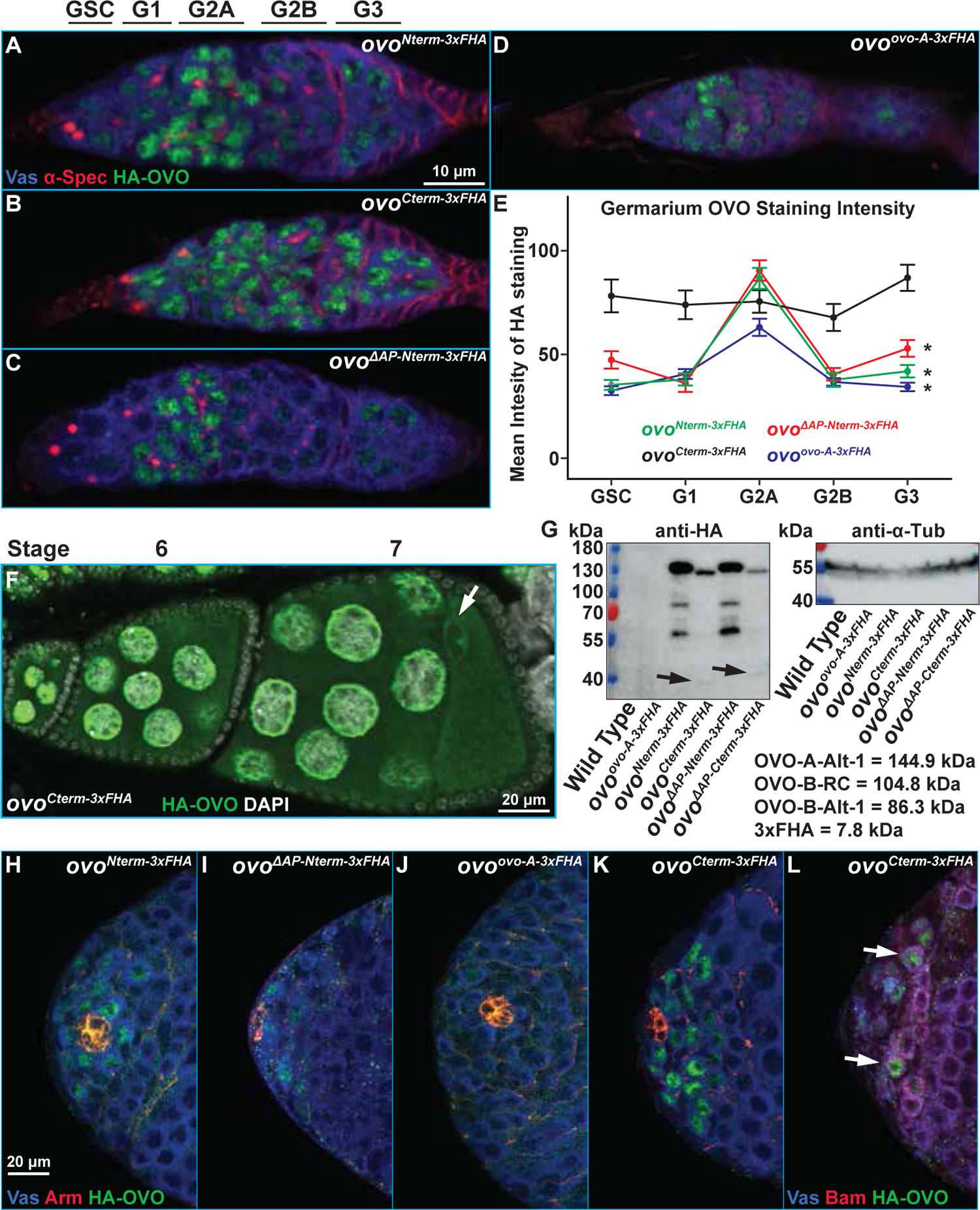
OVO Localization Within the Adult Germline. A-D) Immunofluorescent staining of adult germariums of the indicated genotypes (63x, scale bar = 10 μm). Germariums were stained for Vas (blue) to label the germline, α-Spectrin (red) to label dot spectrosome and fusomes, and HA (green) to label OVO. Germarium regions indicated above A. E) Mean HA-OVO staining intensity for individual nuclei in the indicated germarium regions. Asterisks denote significantly different staining intensities between germarium cell types for each genotype according to a one-way ANOVA test. Error bars represent standard error of the mean. F) Immunofluorescent staining of late stage egg chambers of the indicated genotype. Egg chambers were stained for HA and DAPI. Arrow indicates the oocyte nucleus. G) Western blot for anti-HA and anti-α-Tubulin of the indicated genotypes. Size of the ladder is indicated to the left. H-K) Immunofluorescent staining of adult testes of the indicated genotypes (40x, scale bar = 20 μm). Testes were stained for Vas (blue) to label the germline, Arm (red) to label the somatic cells of the hub, and HA (green) to label OVO. L) Immunofluorescent staining of adult testis of the indicated genotype (40x). Testes were stained for Vas (blue), Bam (red) to label differentiating cysts, and HA (green). Arrow indicates cells that are both Bam and HA positive.

Both ovaries and testes have germ cell niches where germline stem cells self-renew and give rise to daughters that undergo 4 rounds of mitosis with incomplete cytokinesis to create 16-cell cysts. These then begin to differentiate into gametes. Although all female germ cells showed OVO expression, regardless of stage, we noticed differences in anti-HA staining intensity in germ cells in particular germarial stages. The germarium can be divided into the germline stem cells (GSC), G1, G2A, G2B, and G3 regions (Figure 4E). This was especially true for germ cell nuclei in the germarium that were tagged N-terminally (*ovo^Nterm-3xFHA^*, *ovo^ΔAP-Nterm-3xFHA^*, and *ovo^ovo-A-3xFHA^* alleles), which stained the middle region (G2A) significantly more strongly. Oddly, the OVO staining intensity in *ovo^Cterm-3xFHA^* germaria looked more uniform (Figure 4A-D). We decided to measure the normalized mean OVO staining intensity in individual nuclei subdivided by germarium regions in our tagged alleles. We did in fact find that N-terminally tagged *ovo* alleles (*ovo^Nterm-3xFHA^*, *ovo^ΔAP-Nterm-3xFHA^*, and *ovo^ovo-A-3xFHA^*) had significantly higher anti-HA staining intensity in region G2A nuclei compared to the GSC, region G1, G2B, and G3 nuclei, while there were no significant difference in staining intensity between regions in the C-terminally tagged allele *ovo^Cterm-3xFHA^*allele (Figure 4E). This possibly indicated some type of post-translational modification of OVO.

There is no known function for *ovo* in the male germline, but *ovo* reporter genes are weakly expressed in the male germline at the testis apex, where mitotic spermatogonia are found (B. Oliver, Perrimon, and Mahowald 1987; Lü et al. 1998; Bielinska et al. 2005; Andrews and Oliver 2002). Testes from hemizygous *ovo^Nterm-3xFHA^*, *ovo^Cterm-3xFHA^*, and *ovo^ΔAP-Nterm-3xFHA^*males showed anti-HA staining in the spermatogonia (Figure 4H,I,K). This staining was feeble and we had to adjust the laser to very high intensity to detect expression. We were not able to detect anti-HA staining in *ovo^ovo-A-3xFHA^* testes (Figure 4J). Male germline stem cells are directly abutting the somatic niche structure known as the hub. When we co-stained with Arm, which strongly labels the somatic cells of the testes hub, we found that male germline stem cells also expressed OVO-B. Male germ cell differentiation requires Bam (McKearin and Spradling 1990). We also observed OVO-B in Bam positive cells (Figure 4L, arrow). However, not every Bam positive cell, or male germline stem cell in contact with the hub was OVO positive. We do not know if this is stochastic or if OVO expression is near the detection limit of our tools. We never observed OVO in meiotic spermatocytes or sperm. It is possible that regulatory mechanisms exist to ensure OVO levels are low in spermatogonia, but increased *ovo* copy number increases positive autoregulatory activity in spermatogonia without a major effect on spermatocytes (B. Oliver and Pauli 1998a). The *ovo* core promoter that is required for autoregulation is peculiar and spermatocytes have special core promoter binding proteins (M. Hiller et al. 2004; M. A. Hiller et al. 2001; Lu et al. 2020; Li et al. 2009). It is possible that male germ cells do not contain the appropriate transcriptional machinery for *ovo* expression in spermatocytes.

### Is OVO Controlled at the Post-translational Level?

We were puzzled by the fact that C-terminally tagged OVO-A/OVO-B in *ovo^Cterm-3xFHA^* germaria had broad and strong expression in the germarium, given that all N-terminally tagged alleles showed this increase of nuclear mean staining intensity in the middle region of the germarium (increasing from GSC to G2A, followed by a decrease from G2A to G3). The only difference between alleles was the location of the 3xFHA tag. This suggests that OVO might undergo some type of post-translational regulation in the germarium. The 5’ UTR of *ovo* is complex with multiple out-of-frame short ORFs characteristic of translational control (Andrews, Levenson, and Oliver 1998) as well as multiple in-frame initiation codons downstream of the N-terminal tags. It is possible that there is a family of OVO isoforms due to translational control. Another possibility is a proteolytic cleavage event similar to the regulatory mechanism of SVB, where a small peptide, encoded by *tarsal-less*, is involved in directing a proteolytic cleavage of the SVB N-terminus to change the activity of SVB from repressive to active (T. Kondo et al. 2010).

For a first look at possible post-translational regulation at *ovo*, we determined the size of the tagged OVO proteins by western blot. OVO-A and OVO-B are predicted to be 144.9 kDa and 104.8 kDa, respectively, while the addition of the 3xFHA tag would add another 7.8 kDa. We isolated protein from 3-5 day old wild _type, *ovo*_*_ovo-A-3xFHA_*_, *ovo*_*_Nterm-3xFHA_*_, *ovo*_*_ΔAP-Nterm-3xFHA_*_, *ovo*_*_Cterm-3xFHA_*_, and *ovo*_*_ΔAP-Cterm-3xFHA_* _ovaries and loaded_ equal amounts of total protein for gel electrophoresis, membrane transfer and anti-HA staining (Figure 4G). We detected no bands from extracts of HA negative wild type controls or *ovo^ovo-A-3xFHA^* ovaries. Thus, there was no false positive staining observed. These data also suggested that the levels of OVO-A protein are low, which is consistent with transcriptional profiling and ovary staining experiments.

HA positive bands were detected in extracts from *ovo^Nterm-3xFHA^*, *ovo^ΔAP-Nterm-3xFHA^*, *ovo^Cterm-3xFHA^*, and *ovo^ΔAP-Cterm-3xFHA^*ovary samples (Figure 4G). We noticed differences between the apparent band sizes and intensities that were specific to the placement of the 3xFHA tag on OVO, and not specific to whether the allele contained a deletion of the *ovo-A* promoter or not. A prominent band at slightly greater than ∼130 kDa was found in both the *ovo^Nterm-3xFHA^*and *ovo^ΔAP-Nterm-3xFHA^* samples. Although this band was in between the predicted sizes of OVO-A and OVO-B, the fact that we were unable to detect a band in the *ovo^ovo-A-3xFHA^* sample likely indicates that this ∼130 kDa band corresponds to OVO-B. The *ovo^Nterm-3xFHA^* and *ovo^ΔAP-Nterm-3xFHA^*ovary extracts also had smaller prominent HA bands at roughly ∼85 kDa and ∼60 kDa. These smaller N-terminal peptides might be expected from either a cleavage event or translational termination before the zinc-finger domains.

The C-terminally tagged OVO-B extracts had a slightly smaller major band running at ∼120Kda, which could be due to a strong downstream initiation codon at amino acid position 441. There were also faint HA positive bands at ∼40 and ∼80kDa (Figure 2G, arrows). The sum of these products was close to the size of the largest OVO-B HA band. This is consistent with a post-translational cleavage event regulating OVO-B. However, the second largest ∼80 kDa band in *ovo^Nterm-3xFHA^* could also result from OVO-B isoform that does not contain the extended exon 3, which is predicted to be 86.3 kDa. The final OVO-B band at ∼60 kDa is a mystery. Regardless of the mechanism, it appears that OVO proteins outside of the G2A region have different N-termini than those within G2A. While there is clearly much more work to be done, we suggest that examining the translational and post-translational control of *ovo* in the germarium would be worthwhile.

### OVO is maternally deposited and persists in the germline throughout embryogenesis

Regardless of staining differences between N- and C-terminally tagged OVO in the germarium, in later stages of oogenesis, *ovo^Cterm-3xFHA^*showed a persistent nuclear staining in the germline throughout cyst maturation. Nuclear OVO staining was especially prevalent in developing nurse cells (Figure 4F). We also detect OVO staining in the oocyte cytoplasm, with a clearly visible ring of OVO staining around the oocyte nucleus itself (Figure 4F, arrow). The oocyte nucleus is arrested in meiosis and is largely transcriptionally inert with chromosomes highly compacted into a karyosome (Hughes et al. 2018). It would be interesting to know if this karyosome OVO is repressive, activating rare transcribed genes, or simply using the oocyte nucleus to arrive into the mature egg. Staining of the oocyte cytoplasm raises the possibility that OVO is maternally deposited in the embryo. To address this possibility, we stained embryos.

We first looked at 90 minute old embryos derived from *ovo^Cterm-3xFHA^*females, which is around stage 4 of embryogenesis and right before cellularization (Figure 5A). Embryos were immunostained for anti-HA, labeling maternal OVO, and anti-Vas, labeling pole cells. We were primarily interested in assessing the localization and distribution of maternal OVO and reasoned that any positive anti-HA staining should be strictly due to maternal deposition. Although our C-terminal tag alleles will also inherently tag SVB, there is no known role for SVB at this time point in embryonic development or evidence that SVB is zygotically expressed at this time point somatically as well. Also, zygotic transcription does not begin at this stage in pole cells, so any anti-HA staining in the pole cells should be strictly due to maternal deposition (Van Doren, Williamson, and Lehmann 1998). OVO immunostaining of embryos at stage 4 showed what we hypothesize is a progression from evenly distributed non-subcellular specific staining throughout the length of the embryo, to a more nuclear specific localization especially noticeable in the posterior of the embryos (Figure 5B). This transition to nuclear localization progressed to increased nuclear localization in newly formed primordial germ cells (pole cells) and nearby somatic nuclei, to strong localization exclusively in pole cells (Figure 5C). Thus, OVO nuclear localization proceeds pole cell formation and is not a consequence of pole cell formation - a temporal relationship that may be significant.

**Figure 5:**
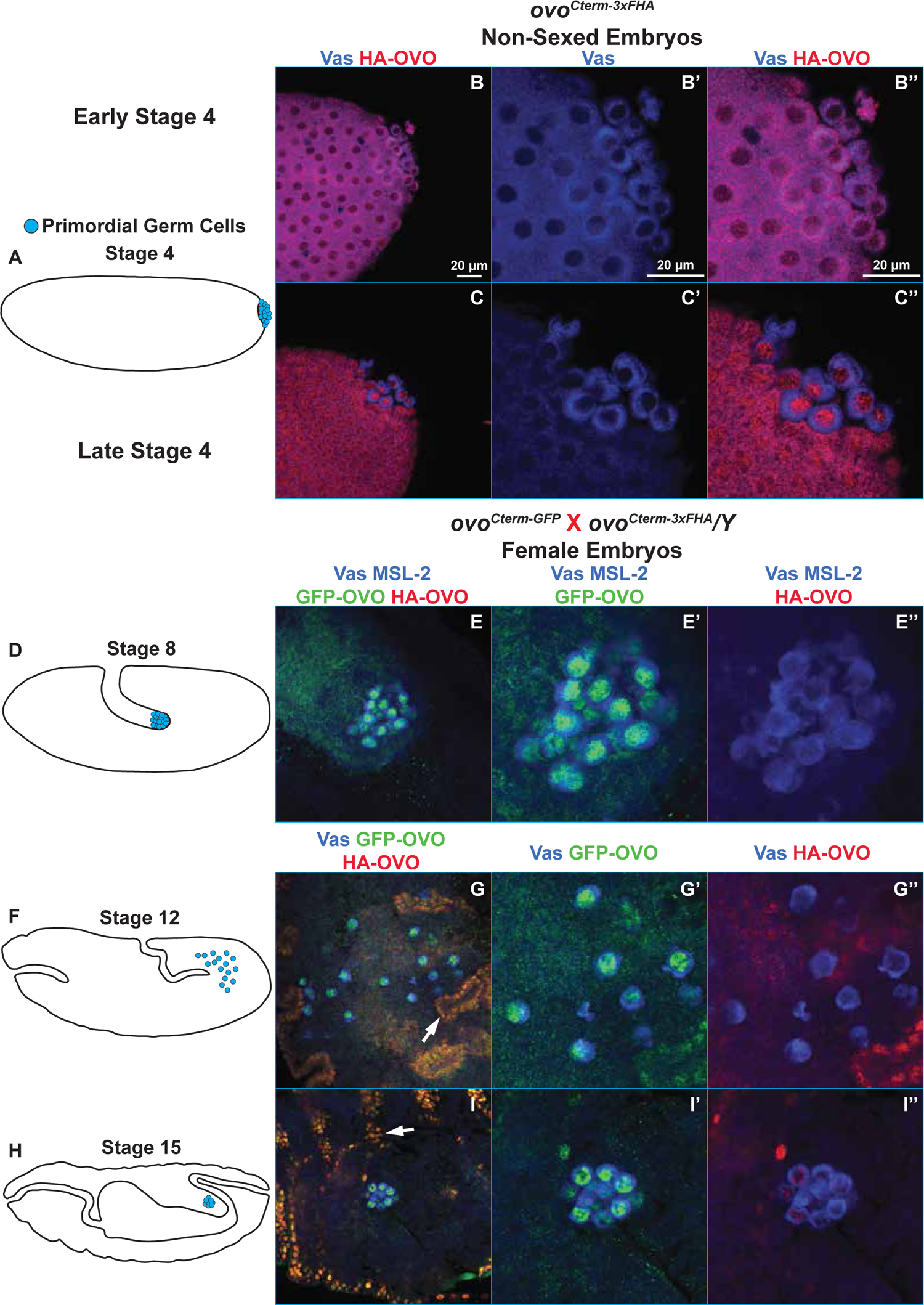
Maternal and Zygotic OVO in Female Embryonic Germ Cells. A,D,F,H) Cartoon of Drosophila embryogenesis stages indicating the location of primordial germ cells with blue dots. B,C) Immunofluorescent staining of embryos derived from *ovo^Cterm-3xFHA^*mothers (40X, scale bar = 20 μm). B, B’, B’’) Early stage 4 embryos were stained for Vas (blue) to label the pole cells and HA (red) to label OVO. C, C’, C’’) Late stage 4 embryos were stained for Vas (blue) and HA (red). E,G,I) Immunofluorescence staining of embryos derived from *ovo^Cterm-GFP^* mothers crossed to *ovo^Cterm-3xFHA^* fathers (40X, scale bar = 20 μm). E, E’, E’’) Stage 8 embryos were stained for Vas (blue), MSL-2 (blue) to indicate male progeny, GFP (green) to label maternal OVO, and HA (red) to label zygotic OVO. Vas and MSL-2 could be differentiated based on their perinuclear vs nuclear puncta staining, respectively. G, G’, G’’) Stage 12 embryos were stained for Vas (blue), GFP (green), and HA (red). Somatic cells at this developmental stage were positive for anti-GFP and anti-HA staining, due to the expression of zygotic SVB, thus allowing us to sex embryos based on positive somatic GFP only, versus positive somatic GFP and HA staining. I, I’, I’’) Stage 15 embryos were stained for Vas (blue), GFP (green), and HA (red). Embryos were sexed based on positive somatic GFP only, versus positive somatic GFP and HA staining. Prime images are zoomed insets of the germ cells. White arrows indicate SVB staining.

Some pole cell localization of maternally derived mRNAs and proteins are transient, while others, such as the pole cell granule protein Vas are persistent. We continued to follow the pattern of OVO localization and expression throughout embryogenesis, as the pole cells invaginate during gastrulation and migrate to the future location of the gonads. For these experiments, we also wanted to distinguish maternal and zygotic OVO. To achieve this, we crossed females bearing an OVO-B-GFP expressing allele (*ovo^Cterm-GFP^*) to males bearing an OVO-B-HA allele (*ovo^Cterm-3xFHA^*). Based on our cross scheme, anti-GFP will detect both maternal and zygotic OVO in males and females, and anti-HA would detect zygotic OVO only in females (*ovo* is X-linked). We also co-stained with anti-Male-specific lethal-2 (MSL-2) antibodies, therefore allowing us to differentiate between male and female embryos since only males should be positive for MSL-2; a component of the somatic dosage compensation machinery (Lyman et al. 1997). At stage 8, which is after the germ cells passively migrate during gastrulation but right before they migrate through the midgut (Figure 5D)(A. C. Santos and Lehmann 2004), there is evidence of early germline transcription (Van Doren, Williamson, and Lehmann 1998). At this stage there was no evidence of somatic OVO (25/25), but strong positive anti-GFP staining in the germline (Figure 5E), suggesting that OVO was protected from turnover in the germline or that zygotic *ovo* expression had begun. We observed no detectable paternally derived OVO-B-HA expression in these germ cells. These data indicate that maternal OVO persists during gastrulation in the female germline.

After stage 8, germ cells traverse the midgut and actively migrate towards and associate with the gonadal mesoderm at stage 12 (Figure 5F)(A. C. Santos and Lehmann 2004). In these stages, we began to observe zygotic expression of SVB-HA (as alluded to earlier, the C-terminus of OVO is shared with SVB), which based on the segregation of X chromosomes indicated that they were female. Males at this stage expressed SVB-GFP in the cuticular pattern previously reported (Figure 5G). Maternal OVO-B-GFP was present (25/25), but we did not observe the diagnostic OVO-B-HA zygotic expression in the germ cells (Figure 5G). This indicated that maternal OVO persisted in all germ cells at this stage and suggested that zygotic transcription of OVO had not yet begun.

After germ cells associate with the gonadal mesoderm, the germ band retracts and germ cells fully coalesce with their somatic counterparts by stage 15 (Figure 5H)(A. C. Santos and Lehmann 2004). We looked at anti-GFP and anti-HA staining in stage 15 embryos and found that the somatic SVB staining patterns from the paternally derived OVO-B-HA that we witnessed at stage 12 were still apparent at stage 15, therefore allowing us to continue to sex embryos by SVB staining. We found that female germ cells at stage 15 were all positive for nuclear anti-GFP staining (25/25). We also began to see evidence of zygotic *ovo* expression, as 48% (12/25) of female embryos showed HA positive germ cell staining (Figure 5I). This suggested that zygotic transcription of *ovo* began at or before stage 15 in female germ cells and importantly, maternal OVO still persisted.

To more carefully assess maternal vs zygotic localization of OVO in male embryos, we used male-specific expression of MSL-2. In the progeny of *ovo^Cterm-GFP^* females crossed to *ovo^Cterm-3xFHA^* males (Figure 6A), all stage 8 MSL-2 positive embryos showed nuclear anti-GFP staining in the germline (25/25), confirming that maternal OVO persisted in the male germline (Figure 6C). At stage 12, embryos could be sexed based on somatic SVB staining. All embryos that were positive for GFP only in the soma showed a persistent nuclear anti-GFP staining in the germline (25/25) indicating the presence of maternal OVO (Figure 6D). The same GFP localization pattern was also found in all stage 15 male embryos (25/25)(Figure 6E). Since zygotic OVO was first detected in female germ cells at stage 15, we reasoned that the persistent GFP staining up until stage 15 in male germ cells was due to maternally deposited OVO. To confirm onset of zygotic expression, we crossed *ovo^Cterm-3xFHA^* males to attached-X (*C(1)DX*) females (Figure 6B), forcing paternal transmission of the tag to sons. We co-stained embryos from this cross with MSL-2 to differentiate males from meta-females (*X/C(1)DX*), and found that the earliest sign of anti-HA staining occurred in stage 15 embryos, with 44% (11/25) of these male embryos having detectable nuclear anti-HA in the germline (Figure 6F). Thus, zygotic expression of *ovo* occurs at the same stage in females and males. Previous *in situ* localization of *ovo* mRNA showed a slight female-biased *ovo* expression at stage 15 (Casper and Van Doren 2009). Interestingly, we did not observe an overt sex-bias in OVO protein expression at this stage.

**Figure 6:**
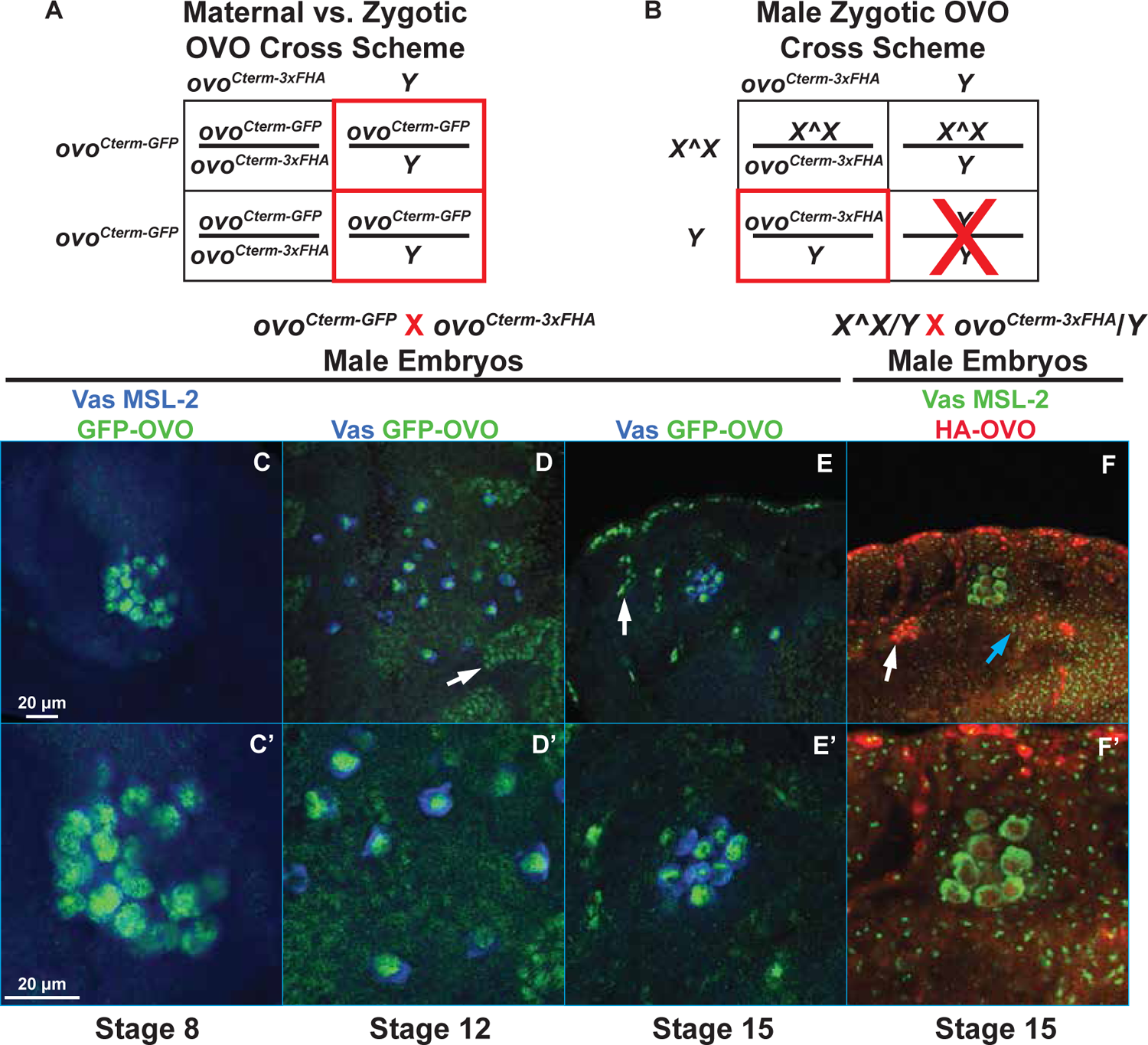
Maternal and Zygotic OVO in Male Embryonic Germ Cells. A,B) Punnett square of the indicated crosses and imaged progeny highlighted with red boxes. C-E) Immunofluorescent staining of embryos derived from *ovo^Cterm-GFP^* mothers crossed to *ovo^Cterm-3xFHA^* fathers (40X, scale bar = 20 μm). Male progeny will inherit the *ovo^Cterm-GFP^* allele as well as maternally deposited C-terminally tagged GFP-OVO. C, C’’) Stage 8 embryos were stained for Vas (blue) to label the germline, MSL-2 (blue) to indicate male progeny, GFP (green) to label maternal OVO, and HA (red) to label zygotic OVO. Vas and MSL-2 could be differentiated based on their perinuclear vs nuclear puncta staining, respectively. D, D’) Stage 12 embryos were stained for Vas (blue), GFP (green), and HA (red). Somatic cells at this developmental stage were positive for anti-GFP and anti-HA staining, due to the expression of SVB, thus allowing us to sex embryos based on positive somatic GFP only, versus positive somatic GFP and HA staining. E, E’) Stage 15 embryos were stained for Vas (blue), GFP (green), and HA (red). Embryos were sexed based on positive somatic GFP only, versus positive somatic GFP and HA staining. F, F’) Stage 15 embryos from *C(1)DX*/*Y* mothers crossed to *ovo^Cterm-3xFHA^*fathers were stained for Vas (green), MSL-2 (green), and HA (red). Male progeny inherit their father’s X chromosome in this cross scheme. Vas and MSL-2 could be differentiated based on their perinuclear vs nuclear puncta staining, respectively. Prime images are zoomed insets of the germ cells. White arrows indicate SVB staining, blue arrows indicate MSL-2 staining.

## Discussion

We find OVO in all adult female germ cells, maternal OVO in the egg and newly formed germ cells, and translation of zygotically expressed *ovo* after activation of germ cell transcription. Maternal deposition persists up until its zygotic expression begins in both males and females (Figure 7). This finding is quite significant given the evidence that OVO positively regulates its own expression (Bielinska et al. 2005; Lü et al. 1998; Andrews et al. 2000). This gap-free expression of positive autoregulatory OVO-B raises the possibility that maternal OVO is necessary for its eventual zygotic expression. While it is technically challenging given the fact that *ovo* is required for egg production, it would be interesting to assess whether removing maternal OVO prevents zygotic *ovo* expression and is required for determining the fate of germ cells without OVO. Since nuclear localization of OVO proceeds pole cell formation, it is possible that the absence of maternal OVO would result in a grandchildless phenotype in both male and female progeny due to failed pole cell formation, or early pole cell loss. Finally, we observed that *ovo* is required during larval stages using a flip-out construct. This potentially indicates that maternal OVO is eventually replaced with zygotic OVO throughout larval germ cell development, and without zygotic *ovo* expression, tested in our flip-out experiments, female germ cells eventually succumb to the absence of OVO and die. We hypothesize that OVO is eternal in the female germline.

**Figure 7.**
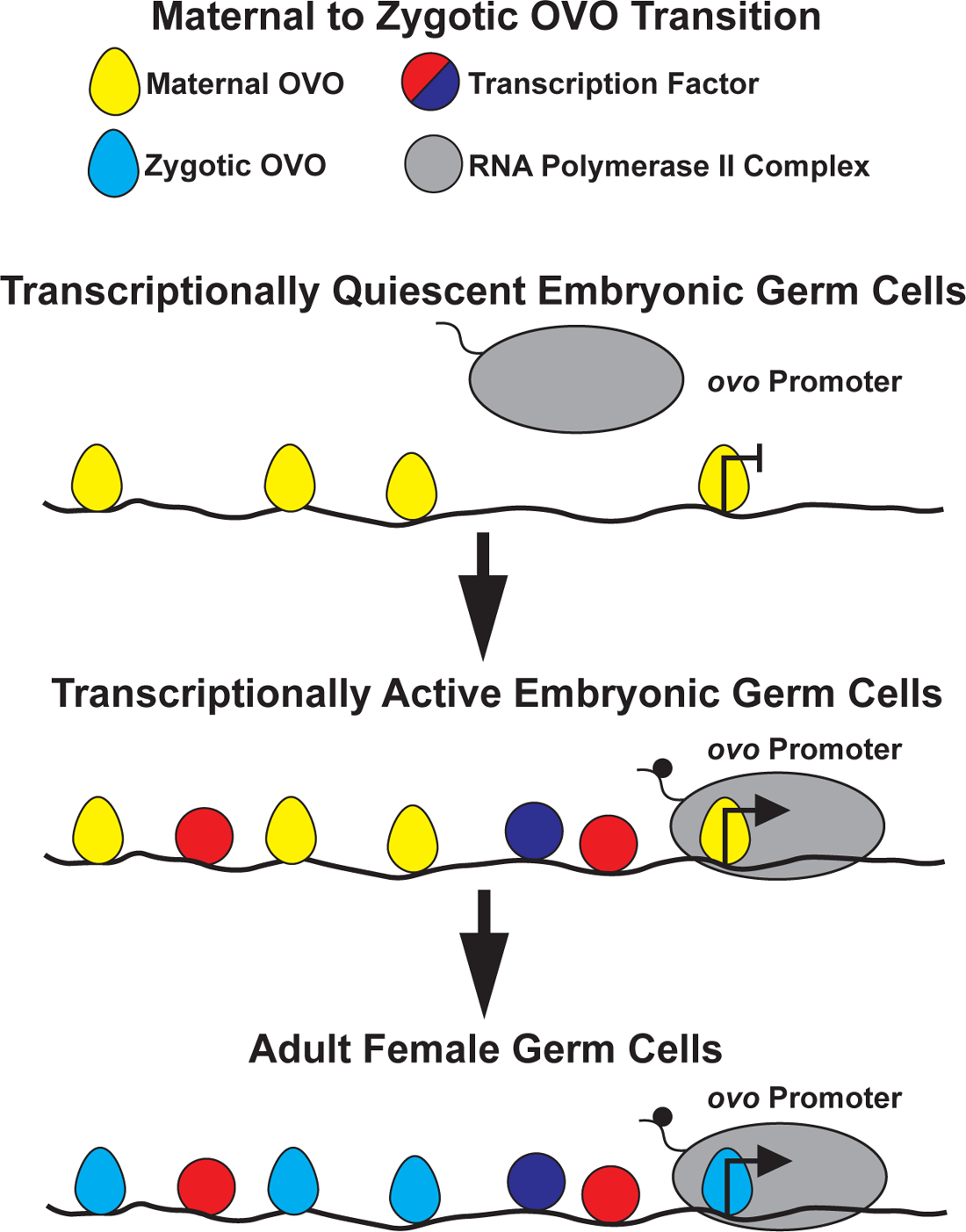
Maternal to Zygotic OVO Transition. Maternally deposited OVO shows nuclear localization in transcriptionally quiescent pole cells during early embryogenesis. This nuclear localization pattern of maternal OVO is persistent throughout embryogenesis in the developing germline and continues until zygotic OVO localization is first detected, even though this is well after the embryonic germline begins zygotic transcription. Zygotic OVO presumably replaces maternal OVO throughout larval germline development and into the adult stages. *ovo* exists in a positive autoregulatory loop and it is plausible that maternal OVO is required for zygotic *ovo* expression, thus ensuring an eternal presence of OVO in the female germline.

While there is a great deal of dated literature on the *ovo* locus, recent technical advances provide an opportunity to confirm and extend those studies. Perhaps, most significantly, we now have clean null alleles of *ovo-*B, rather than the legacy alleles that our data suggests are recessive antimorphs. Design of these null alleles was greatly aided by RNA-seq informed reannotation of the locus. Additionally, we now have a T2A-GAL4 tool to drive expression in cells that are translating OVO and a collection of *UAS-ovo* cDNAs for rescue and overexpression work. Previous work using transgenes at uncontrolled, ectopic genome locations, showed that *ovo* is strongly and broadly expressed in the female germline and weakly expressed in male germline stem cells and spermatogonia (Lü and Oliver 2001; Bielinska et al. 2005; Lü et al. 1998; B. Oliver et al. 1994; Andrews and Oliver 2002). We now know that functional endogenous OVO is expressed in the same pattern.

Previously, we thought that there was a single *ovo* function, with variance in the phenotypes of the five dominant negatives, two key *ovo^D1^*mutations, and weaker hypomorphs from the Mohler female sterile screen being due to quantitative differences (Mohler and Carroll 1984; Andrews, Levenson, and Oliver 1998; Mével-Ninio et al. 1996; Busson et al. 1983; B. Oliver, Perrimon, and Mahowald 1987). It now seems likely that there are distinct OVO functions of the two OVO-B isoforms encoded by exon 2 splice variants. Like SVB, which undergoes a critical regulated proteolytic cleavage to generate isoforms with different functions (T. Kondo et al. 2010), we find that there are different OVO isoforms visualized by N- and C-terminal tags. We do not know if this is a translational event (e.g. due to alternative in-frame AUGs) or proteolytic cleavage, but we do note that the sum of some of the smaller OVO isoforms conveniently approach the apparent size of the longest isoforms. Regardless of mechanism, the N-terminal containing OVO-B isoforms are most abundant in the middle region of the germarium (region 2A). The 16-cell germline cyst is fully formed and the oocyte is specified at this point, and the egg chambers will acquire follicle cells and begin the dramatic transformation leading to an egg.

The function of OVO-A has been studied in the dominant *ovo^D^*alleles, where it is clear that OVO-A is a direct negative autoregulator of both *ovo* promoters, as well as a direct negative regulator of the *otu* target gene. The function of the wildtype OVO-A isoform has never been clear and our work does not shed any further light. Previously, our lab saw that *ovo^D1rv^* females rescued with an OVO-B encoding transgene (*P{ovo^ΔAP^}*) were fertile with a semi-penetrant and ephemeral grandchildless phenotype, leading us to propose that OVO-A might be required to negatively regulate OVO-B production (Andrews and Oliver 2002). Specifically, the hypothesis was that since *ovo-B* exists in an autoregulatory loop, the presence of OVO-A was important to dampen the expression of the feed-forward activity of OVO-B, therefore ensuring that the female germline doesn’t express detrimental or toxic levels of OVO-B. There are two lines of evidence to suggest that this is wrong. First, we showed that maternal OVO-B is abundant and localized to early germ cells, and that zygotic OVO-B begins to accumulate before the maternal supply is depleted. Maternal OVO-A is thus unlikely to be required to turn OVO-B off in the embryo, although it might help prevent runaway autoregulation. Secondly, and most importantly, we found that 98% of daughters from *ovo^ΔAP^* mothers had a germline. This does not mean that the transition from *ovo* regulation of oogenesis, to possible regulation of a completely different primordial germ cell role is uninteresting. It does mean that the role of OVO-A is a mystery. Indeed, it is possible that low level OVO-A simply does no harm and is tolerated. The *ovo-A* ORF shares coding sequences with *svb* and uses a common splice acceptor (5’ of exon 2), so it is entirely possible that *ovo-A* is effectively a leaky, but innocuous, expression of *svb*.

Female and male germ cells in stage 15 embryos both expressed zygotic OVO, however, only females showed a requirement for OVO and only females have abundant OVO in adult germ cells. We carefully sexed embryos using both differently tagged *ovo* alleles and staining with antibodies against the male specific somatic dosage compensation complex. Previous work showed that *ovo* mRNA was more abundant in female embryos than male embryos in the germline (Casper and Van Doren 2009). This is certainly logical, given the female-specific requirement for OVO, but we did not observe female-biased OVO protein levels in our work here. Given the autoregulation of *ovo*, and a presumably equal maternal load of OVO in the primordial germline, any regulated decrease in *ovo* expression would likely require another gene activity besides *ovo* itself. It has been previously shown that the *ovo* core promoter incubated in either ovary or testes protein extract show differences in mobility in gel-shift assays (Bielinska et al. 2005). This finding suggests that the repertoire of proteins binding to the *ovo* core promoter differ among the sexes. It has not escaped our attention that *ovo* is X-linked and therefore females have two doses and males have one. Regardless, it would be interesting to determine how exactly the male germline eventually reduces and diminishes *ovo* expression in stem cells and spermatogonia, and what turns off *ovo* more completely during the shift from spermatogonia to spermatocyte. Discovering the difference in OVO regulation between male and female germ cells will offer further insight into the establishment of sexually dimorphic gene expression and cellular identity leading to sex-specific gamete production.

## Data Availability

Drosophila strains and plasmids used for this study are available upon request. All sequence information and datasets used in this study are in Table S1.

## Acknowledgements

We would like to thank previous and current members of the Oliver lab, Laboratory of Biochemistry and Genetics at NIH, and L.B. committee members Mark Van Doren and Allan Spradling for insightful discussion and comments on this work throughout. Stocks obtained from the Bloomington *Drosophila* Stock Center (National Institutes of Health (NIH) grant P40OD018537) were used in this study. Monoclonal antibodies were obtained from the Developmental Studies Hybridoma Bank, created by the Eunice Kennedy Shriver National Institute of Child Health and Human Development (NICHD) of the NIH and maintained at the Department of Biology, University of Iowa, Iowa City, IA 52242. Genetic and genomic information was obtained from FlyBase (U41 HG-000739). PacBio long-read sequencing was completed by the NIH Intramural Sequencing Center. This work utilized the computational resources of the NIH High-Performance Computing Biowulf cluster (http://hpc.nih.gov). LAB acknowledgements

---

**Table S1:** FlyBase ART Table

**Table S2:** Phenotype Scores for *ovo* Alleles

**Table S3:** PacBio Long-Read Transcripts in GTF Format

**Table S4:** Phenotype Scores for *ovo* Promoter Deletion Alleles

